# Developmental analysis of the cone photoreceptor-less little skate retina reveals distinct Onecut1 isoforms

**DOI:** 10.1101/2025.07.30.667746

**Authors:** Chetan C. Rangachar, Denice D. Moran, Mark M. Emerson

**Affiliations:** Department of Biology, The City College of New York, City University of New York, New York, NY 10031, USA; Biology PhD Program, Graduate Center, City University of New York, New York, NY, 10016; Biochemistry PhD Program, Graduate Center, City University of New York, New York, NY, 10016

**Keywords:** Retina, Photoreceptors, Rods, Cones, Skate, Development

## Abstract

The retinal development of elasmobranchs, the subclass comprising sharks, skates, and rays, remains poorly understood. This group is diverse in retinal phenotype, with many sharks and rays possessing rods together with one or more cone types. In contrast, the little skate (*Leucoraja erinacea*) has only a single rod photoreceptor type, which has been reported to exhibit some physiological and anatomical properties associated with cones. To investigate how this unusual photoreceptor system develops, we first identified an embryonic stage of early photoreceptor formation based on *otx2* expression. We then developed a retinal electroporation approach to test whether a *onecut1*-dependent cone-associated reporter could be activated in the embryonic skate retina. Activation of this reporter was not detected, indicating that the corresponding enhancer is not robustly active under the conditions tested. To assess developmental changes in gene expression, we generated bulk RNA-seq datasets from embryonic, hatchling, and adult retinas. These analyses showed strong embryonic expression of *onecut1*, increasing expression of rod-associated genes through development, and pseudogenization or loss of multiple cone-enriched genes. We further identified a developmentally regulated *onecut1* splice isoform containing an additional 48 amino acid sequence between the CUT and homeodomain DNA-binding domains. This spacer-containing isoform, termed LSOC1X2, was most abundant in the embryonic retina. To test whether LSOC1X2 retained regulatory activity, we assayed it in a mouse retinal reporter system. Both skate Onecut1 isoforms activated the ThrbCRM1 reporter in this heterologous context. Together, these findings identify a novel, developmentally regulated retinal *onecut1* isoform in the little skate and establish it as a candidate regulator for future studies of photoreceptor development in this species and its elasmobranch relatives.

## Introduction

### Evolutionary Conservation of Photoreceptor Subtypes in the Vertebrate Retina

The vertebrate retina exhibits a striking degree of structural conservation across the subphylum. A recent meta-analysis of single-cell retinal transcriptomes across diverse vertebrates found that the major retinal cell classes are broadly conserved, with much of the observed variation arising within those classes rather than between them^1^. In the mature vertebrate retina, six neuronal classes are classically recognized: rod photoreceptors, cone photoreceptors, bipolar cells, horizontal cells, amacrine cells, and retinal ganglion cells. These are accompanied by a single resident glial class, the Müller glia^2^.

Chondrichthyans, or cartilaginous fishes, represent an important comparative group for understanding the evolution of vertebrate retinal development. This lineage diverged from the osteichthyan lineage leading to humans roughly 459 million years ago^3^, yet remains relatively understudied in developmental retinal biology. Extant chondrichthyans comprise two major groups: holocephalans and elasmobranchs. Holocephalans include the chimaeras, whereas elasmobranchs include sharks, skates, and rays^4^. Phylogenetic analysis of vertebrate opsins suggests that the ancestral vertebrate photoreceptor was cone-like, with rods arising later as a specialized photoreceptor type^5^. A recent nomenclature reflecting this evolutionary framework has proposed referring to rods as PR0 and cones as PR1–PR6^6^. Under this framework, the chimera Australian ghost shark (*Callorhinchus milii*) possesses PR1 and PR2 photoreceptors, suggesting that the ancestral chondrichthyan retina contained multiple cone classes^7^. Among extant elasmobranchs, however, photoreceptor complements are variable. Many sharks retain a single cone type, usually PR1 or PR2, while many stingrays possess both PR1 and PR2 (reviewed in ^8^). The little skate (*Leucoraja erinacea*), by contrast, appears to lack cones entirely and instead possesses only a single rod photoreceptor type^9^. Although the skate rod has some unusual functional properties relative to rods in other vertebrates, its transcription factor profile, phototransduction machinery, including rhodopsin, and cellular morphology support its designation as a PR0 rod. This raises a central question: how was the cone photoreceptor fate lost in the skate lineage?

The little skate is especially interesting because its single rod type combines canonical rod features with unusual light-adaptive behavior. Like other rods, it is capable of single-photon responses in the dark-adapted state. However, after adaptation to bright backgrounds, skate rods accelerate their response kinetics and support photopic vision, giving them both a rod-like and a cone-like operating mode^10^. This form of adaptation appears unusual among well-studied vertebrate models and has also been described in the winter skate ^11^. Ultrastructural studies further suggest that the skate rod synapse shares some features with cone synapses, including large invaginations at the tripartite synapse, multiple ribbons within single invaginations, and long telodendria^12^. Together, these observations suggest that the skate visual system did not simply eliminate cones and leave rods unchanged. Instead, cone-associated features may have been lost, retained, or repurposed in distinct ways. How these evolutionary changes are reflected in retinal development remains unclear.

One candidate gene of particular interest in this context is *onecut1*, a transcription factor with a well-established role in cone, horizontal, and retinal ganglion cell development in other vertebrates ^13–16^. If the little skate lost the cone fate through modification of an otherwise conserved developmental program, changes in the regulation, structure, or activity of *onecut1* could provide one molecular window into that transition.

In this study, we examined developmental features of little skate retinogenesis and asked how *onecut1* is deployed in a cone-less retina. We first used histological analysis of the embryonic retina, including Otx2 expression and cell-cycle markers, to approximate the period of photoreceptor production. Guided by this developmental framework, we generated bulk RNA-sequencing datasets from embryonic, hatchling, and adult retina to identify transcriptional changes associated with photoreceptor differentiation. These data revealed initially high but declining expression of *onecut1*, alongside increasing expression of rod-associated developmental markers over time. Rod phototransduction genes increased from embryogenesis to hatching and remained highly expressed in adulthood, whereas examined cone phototransduction genes were not detectably expressed and several showed evidence of mutational decay. We then identified a developmentally regulated alternative splice isoform of skate *onecut1*. Finally, we assessed the potential consequences of this splicing event by testing whether skate Onecut1 isoforms could activate a well-characterized retinal enhancer in explant assays, and by using AlphaFold3 to model how the canonical and spacer-containing isoforms might differ in their interactions with DNA and cofactors.

Our results identify a novel *onecut1* isoform in the little skate and show that both skate Onecut1 isoforms retain the ability to activate a cone-associated enhancer in a heterologous retinal context. Rather than providing a complete explanation for cone loss in skate, these findings identify a specific alteration in a major photoreceptor regulatory factor and establish a tractable molecular entry point for studying how the cone developmental program was modified in a rod-only vertebrate retina.

## Results

### Cell Cycle Dynamics of the Little Skate Retina

We initiated this study by examining the embryo stages of the little skate to determine the early stages of photoreceptor formation and to roughly characterize the cell cycle dynamics at this time, primarily to inform the implementation of electroporation. Because little skate embryos develop over approximately 20 weeks in waters no warmer than 20°C, we reasoned that retinal cell-cycle progression might be slower than in warm-blooded or artificially incubated vertebrate models such as mouse and chick. We chose to examine stage 29 (“S29” ∼50 days post oviposition) skate embryos according to a reported little skate staging system and a developmental staging analysis of the related winter skate which narrowed the window of eye development^17,18^. A preliminary assessment at S29 of Otx2 expression, an early photoreceptor-associated marker, identified that a number of Otx2+ cells were formed suggesting that photoreceptor genesis had begun by this stage ^19^. We injected live S29 little skate embryo yolks with thymidine analog 5’-ethynyl-2’-deoxyuridine (EdU), exposing several groups of specimens to the label for different lengths of time. Embryos were then fixed and analyzed for incorporation of EdU within the phosphohistone H3-positive (PHH3+) mitotic population at each time point. Initial choices of incubation window were broad, from 2 to 48 hours (Fig S1 B,C). However, it was observed that at 4 hours, almost none of the PHH3+ population is EdU+ (1/87 counted cells across four specimens) (Fig 1A, B), and by 8 hours, over 70% of the PHH3+ population is EdU+ (43/56 counted cells across four specimens) (Fig 1C, D). Numerical comparison of the dividing cell population shows a significant increase in EdU uptake between 4 and 8 hours post injection (Fig 1E, S1A). Cells sensitive to uptake of EdU are within the S-phase and begin to express PHH3 as mitosis begins. The shortest time to incorporate the label in the mitotic population is a conservative estimate of the length of G2, the period between the end of DNA synthesis and cell division. More elaborate experiments have been performed in the P1 mouse with a similar methodology, using tritiated thymidine as the DNA replication marker and visual inspection of micrographs to label cells undergoing mitosis. Estimates from this experiment reveal that in the P1 mouse, the average length of G2 is 2.6 hours, and the whole cell cycle is 30 hours in length^20^. If the proportional relationship between G2 length and total cell-cycle length is broadly similar to that reported in the P1 mouse retina, this would place the total cycle length of dividing cells in the embryonic skate retina at roughly 46 hours. We treat this as a coarse approximation rather than a direct measurement. This time frame suggests that a three-day culture should be more than sufficient to read out reporter activity of exogenous plasmids, which are believed to be incorporated into the nucleus after the breakdown of the nuclear envelope in prophase of mitosis.

**FIGURE 1.**
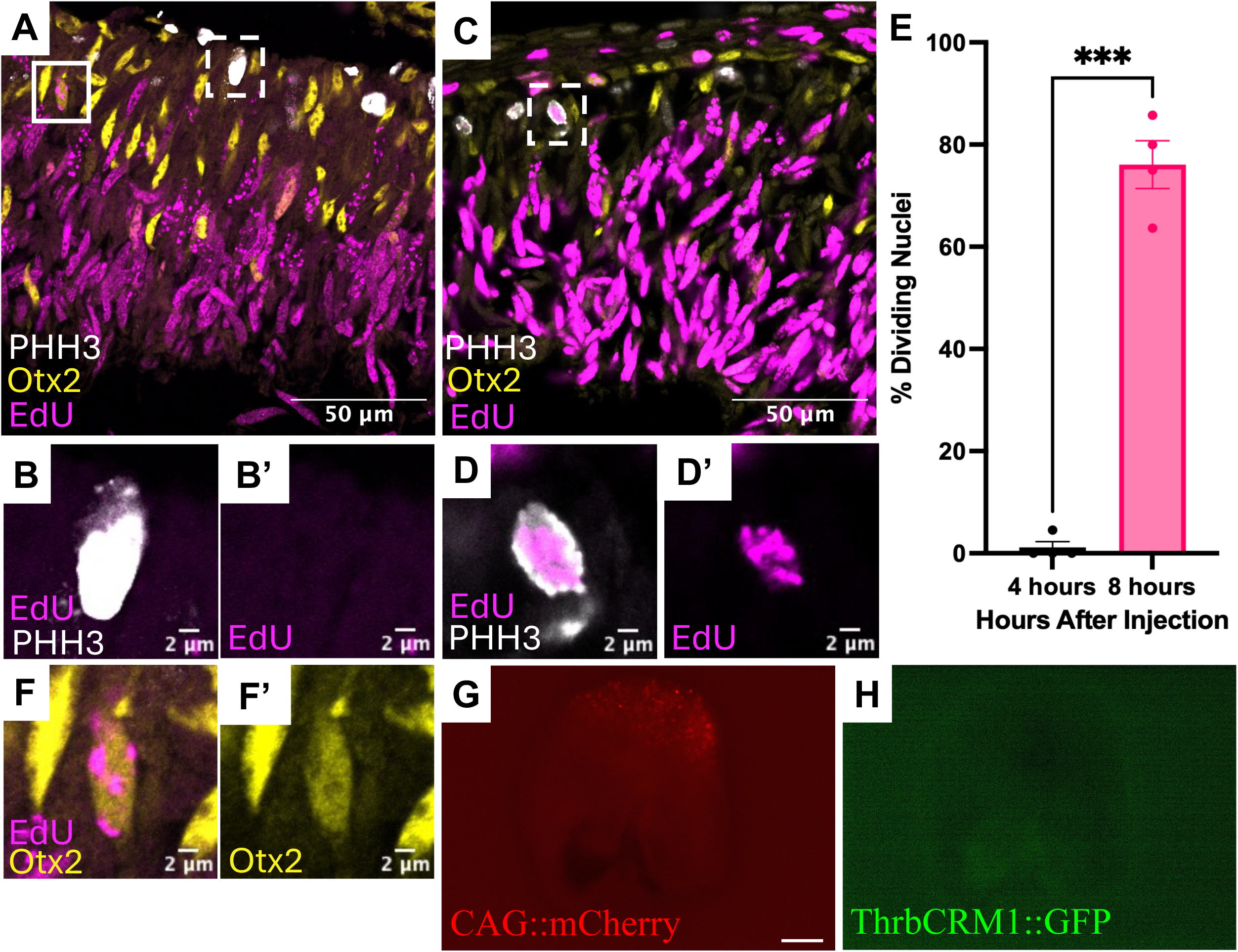
Thymidine birthdating analysis of embryonic little skate retina and electroporation of THRB CRM1 reporter. (A) Embryonic (S29) retina pulse labeled with EdU for 4 hours, stained for EdU in magenta (Alexa Fluor 647), Otx2 in yellow (Alexa Fluor 488), and phospho-histone H3 (PHH3) in white (Cy3). Solid white box indicates representative Otx2+/EdU+ nucleus. Dashed white box indicates representative EdU-/PHH3+ labeled nucleus. (B) Magnified image of representative EdU-/PHH3+ nucleus, with both channels (EdU, PHH3) visualized. (B’) B image with PHH3 channel removed. (C) 8-hour labeled S29 retina. Dashed box indicates representative EdU+/PHH3+ nucleus. (D) Magnified image of PHH3+/EdU+ nucleus with both channels (EdU, PHH3) present. (D’) D image with PHH3 channel removed. (E) Bar graph showing percent of dividing nuclei (PHH3+) with EdU labeling at the time of retinal fixation, by incubation length. p < 0.001 by Welch’s t-test. (F) Magnified image of Otx2+/EdU+ nucleus. (F’) F image with Otx2 channel removed. (G,H) S29 little skate retina electroporated with CAG::mCherry and ThrbCRM1::GFP and visualized in wholemount for mCherry (G) and GFP (H). Scale bar in G = 200 µm and applies to H.

In other vertebrates, such as the mouse and chick, photoreceptor-competent restricted retinal progenitor cells express Otx2, so we analyzed the presence of this developmental marker in the little skate’s dividing (EdU+) population^16^. After 4 hours of EdU exposure, a subset of EdU+ cells also expressed Otx2, consistent with the possibility that photoreceptor-producing progenitors are present in the S29 skate retina (Fig 1F). In other vertebrates, co-expression of Otx2 and Onecut1 in early restricted retinal progenitor cells associated with cone formation activates the cis-regulatory element ThrbCRM^16^. At more mature developmental stages, Onecut1 is no longer expressed in RPCs and rods, but not cones, are formed during this period^14,16^. A significant outstanding question then is whether or not the skate’s Otx2+ dividing population expresses *onecut1*, or if loss of *onecut1* expression is a causal driver of the loss of cones in the little skate. We attempted to detect Onecut1 protein immunohistochemically using a monoclonal antibody raised against mouse Onecut1 (sc-376167, Santa Cruz Biotechnology), but did not obtain interpretable signal, likely because the antibody recognizes an N-terminal epitope that is less conserved across species than the DNA-binding domains.

To test whether an Otx2/Onecut1-like regulatory state could be detected functionally in the embryonic skate retina, we developed an electroporation-based reporter assay by adopting parameters developed for the ascidian *Ciona intestinalis*^21^. To test the electroporation procedure, retinas from S29 embryos were electroporated with GFP and mCherry fluorescent reporters, both driven by the broadly active CAG element^22^. Whole-mount visualization of the retina showed broad co-expression of the two fluorescent reporters, confirmed by confocal analysis of sections (Fig S2 A-E). The ThrbCRM1 element is co-regulated by Onecut1 and Otx2 and evidence suggests that both factors are required for reporter expression^16,23^. We thus co-electroporated ThrbCRM1::GFP reporter with CAG::mCherry, cultured for three days, and assessed the reporters. Although the electroporated region was clearly marked by mCherry, we did not detect GFP driven by ThrbCRM1 under these conditions (N=3 retina) (Fig 1 G,H). This result indicates that the ThrbCRM1 enhancer was not detectably activated in the S29 skate retina but this by itself does not distinguish between absence of an Otx2/Onecut1-like regulatory state and broader species-specific differences in enhancer interpretation or developmental timing.

Together, these findings identify the S29 little skate retina as a tractable developmental stage for testing hypotheses about early photoreceptor specification in a cone-less retina. Successful retinal electroporation establishes an experimental platform for such studies. At the same time, the lack of ThrbCRM1 activation should be interpreted cautiously, as it may reflect failure to target the relevant progenitor population, differences in cell-cycle timing within that population that limit plasmid uptake or features of the endogenous skate regulatory environment that prevent reporter activation. It may also indicate that Otx2 and Onecut1 are not co-expressed in the same cells, or that the ThrbCRM1 enhancer is not interpreted equivalently in skate.

### Bulk RNA-seq Reveals Robust Onecut1 Expression in Early Development, with Rod Program Genes Increasing in Expression in Later Development and Maturity

To examine gene expression broadly across little skate retinal development, we conducted bulk RNA sequencing of the whole retina. Eight specimens across three developmental stages—three S29 embryos, three hatchlings (P0), and two adults—were analyzed. Expression values were normalized as transcripts per million (TPM), log2-transformed, and centered by each gene’s mean expression across stages to visualize changes in developmental transcription factor (Fig. 2A) and phototransduction gene expression (Fig. 2B). Genes in both heatmaps are annotated by functional pathway. Genes in the developmental heatmap were selected to represent transcription factors with established roles in photoreceptor fate induction, guided by the framework reviewed by Wang and Cepko^24^. In the phototransduction heatmap, genes are classified following the framework summarized by Lamb^25^.

**FIGURE 2.**
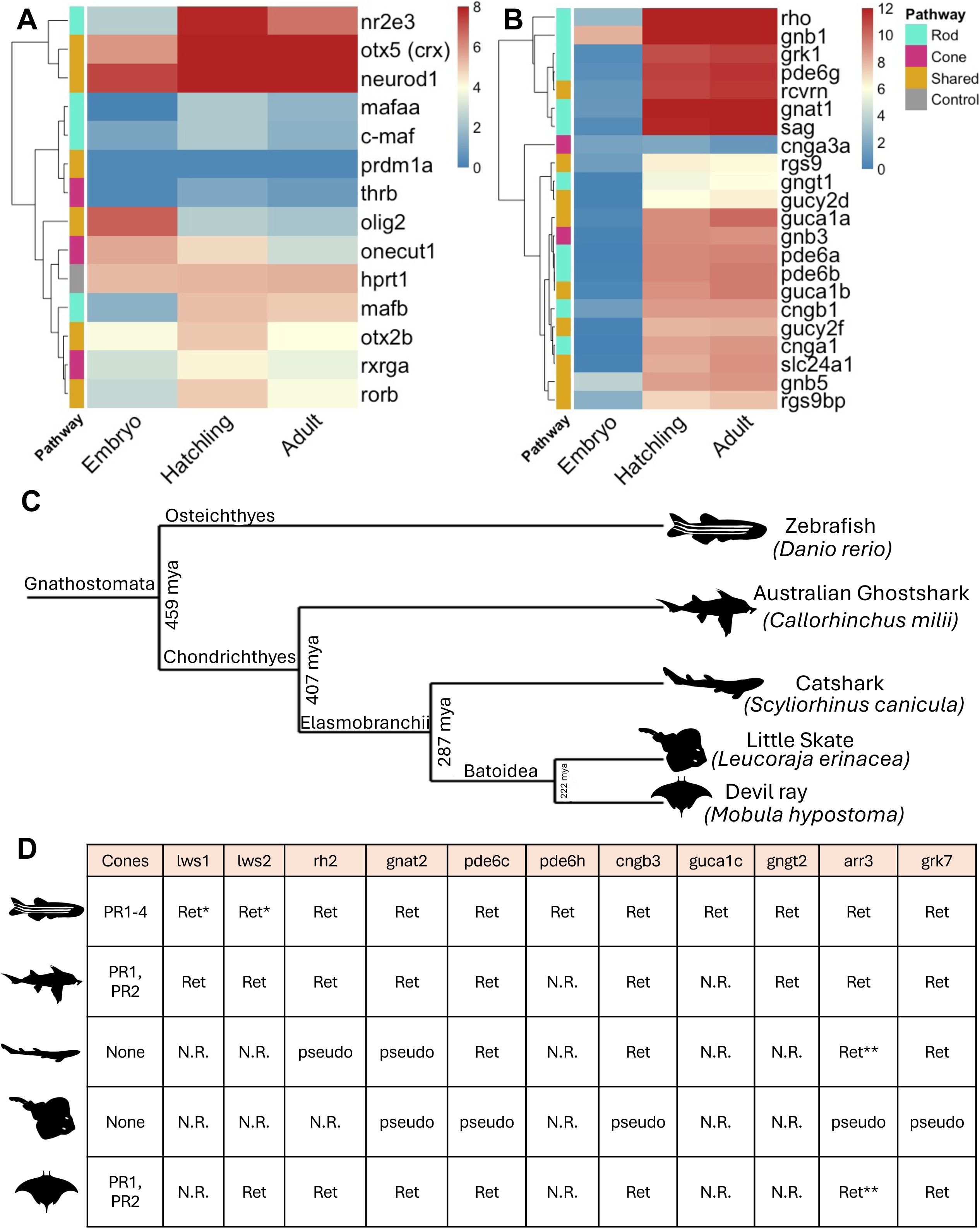
Developmental gene expression profiles of photoreceptor expressed genes and evidence of pseudogenization of cone phototransduction components across Chondrichthyans. (A) Heatmap showing log2-transformed average TPM [log2(TPM + 1)] for transcription factors associated with retinal development across three developmental stages: stage 39 (n=3), hatchling P0 (n=3), and adult (n=2). Color scale spans 0–8 log2 TPM, from steelblue (low) through light yellow to firebrick (high). Color intensity reflects absolute expression level. Genes are clustered by hierarchical clustering using Euclidean distance with complete linkage. The housekeeping gene hprt1 shows minimal variation across stages, as expected for a constitutively expressed control. Genes are grouped by pathway (rod, cyan; cone, magenta; shared, yellow). (B) Heatmap showing average phototransduction gene expression across three developmental stages: S29 embryo (n = 3), hatchling P0 (n = 3), and adult (n = 2). Expression values are shown as log2(TPM + 1). The color scale spans 0–12 log2 TPM. Genes were hierarchically clustered using Euclidean distance and complete linkage. Genes are classified as rod-specific, cone-specific, or shared components of the phototransduction cascade based on ^25^. gnb3 is classified as a cone phototransduction gene in that framework, but has also been characterized as a component of rod bipolar cell signaling conserved across vertebrates^40^. (C) Phylogenetic tree showing the relationships among the Australian ghost shark (Callorhinchus milii), catshark (Scyliorhinus canicula), devil ray (Mobula hypostoma), and little skate (Leucoraja erinacea), with zebrafish (Danio rerio) shown as an osteichthyan outgroup. Divergence times at major nodes are indicated in millions of years ago (mya), based on Marletaz et al.^3^. (D) Table summarizing the genomic status of cone-associated phototransduction genes in the Australian ghost shark, catshark, little skate, and devil ray, shown in the same phylogenetic order as in Fig. 2C. The leftmost column indicates the cone photoreceptor complement reported or inferred for each species. Gene status was assigned on a gene-by-gene basis using genome annotation together with TBLASTN homology searches and syntenic context where informative. Ret., retained intact ortholog; pseudo, pseudogenized locus or remnant identified; N.R., no recognizable ortholog recovered in the current genome assembly. For *arr3, gnat2, pde6c, cngb3,* and *gngt2*, assignments were informed by syntenic comparison where possible. For genes lacking informative syntenic anchors, including *pde6h, guca1c,* and the opsins, assignments were based on whole-genome TBLASTN searches and available annotation. *Zebrafish has annotated long wavelength-sensitive opsins classified as *lws1* (*opn1lw1*) and *lws2* (*opn1lw2*) that arose from a different gene duplication event than the ghost shark genes. **The syntenic context of ghost shark *arr3* was conserved in catshark and devil ray, but at this position each genome contains a gene annotated as “beta arrestin 1-like,” which shows significant alignment to ghost shark *arr3*.

A question of interest in skate photoreceptor development is which large Maf transcription factors are present to support rod-associated developmental regulation. This family includes four orthology classes: MafA, MafB, c-Maf, and NRL^26^. In mammals and in teleost fish, *Nrl* is expressed to drive the rod fate, but the teleost zebrafish has an alternative rod-specifying pathway utilizing *mafba*, one of two copies of MafB present in the zebrafish genome^27^. In avian species such as the chick, *NRL* is absent from the genome, and MafA is thought to serve this role^28^. We looked for all of the Maf family genes in the skate retinal transcriptome, finding only one annotated Maf member, *mafaa*. However, the skate genome is not yet completely annotated, and many relevant genes are labeled by locus numbers (LOC#######). This motivated us to perform TBLASTN analysis using human and zebrafish NRL proteins as queries against the skate genome to determine whether additional large Maf homologs could be identified in this way. We found that zebrafish NRL is most similar to a gene labeled LOC129705330, described as “transcription factor *maf*-like” (73% identity, E = 1e-32), and that human NRL is most similar to a gene labeled LOC129707262, described as “transcription factor *mafb*-like” (39% identity, E = 1e-31) (Fig. S3A). Synteny analysis of the surrounding gene neighborhoods indicates that the *maf*-like gene corresponds to *c-maf*, whereas the *mafb*-like gene corresponds to *mafb*. Less significant alignments were also recovered for the small Maf genes *mafk* and *mafg*, attributable to their shared bZIP domains. This supports the conclusion that a distinct NRL ortholog is absent from the current little skate genome assembly. Of the three large Maf family members identified, *mafb* is expressed at the highest level at each developmental stage, supporting its candidacy as the predominant large Maf factor associated with rod development in the skate retina. Consistent with this interpretation, rod-associated developmental regulators showed a trajectory matching the rod-dominated identity of the little skate retina (Fig. 2A). *mafb* and *nr2e3*, together with the pan-photoreceptor markers *otx5/crx* and *neurod1*, increased in expression from the embryonic stage to hatching and remained elevated in the adult. This pattern is consistent with increased rod photoreceptor differentiation between S29 and hatching.

Against this rod-associated developmental trajectory, *onecut1*, a transcription factor associated with cone specification in other vertebrates, exhibited its highest expression in the S29 embryo and declined through the hatchling and adult stages (Fig. 2A). This pattern is notable given the established role of Onecut1 in early cone-associated developmental programs in other vertebrates^16^. However, other genes associated with this program, including *thrb* and *rxrga*, did not show developmental trajectories that clearly paralleled the embryonic enrichment of Onecut1. In other vertebrates, *Thrb* and *Rxrg* can be driven by the combined activity of Onecut1 and Otx2, but the bulk RNA-seq data presented here do not resolve whether these factors are expressed in the same cells in the embryonic skate retina^16^. Thus, the data indicate that *onecut1* is developmentally enriched in the embryonic skate retina, but do not by themselves establish the cellular context or downstream regulatory consequences of that expression.

The developmental trajectory of mature phototransduction genes further underscores the rod-only identity of the little skate retina (Fig. 2B). Rod-specific phototransduction components, including rhodopsin (*rh1*), rod transducin (*gnat1*), rod arrestin (*sag*), rod phosphodiesterase subunits (*pde6a* and *pde6b*), and the rod cyclic nucleotide-gated channel (*cnga1*), all showed markedly increased expression from embryo to hatchling, with sustained high expression in the adult (Fig. 2B). This pattern is consistent with the increased differentiation and maturation of rod photoreceptors between S29 and hatching.

To place these findings in evolutionary context, we compared the status of cone-associated phototransduction genes across four chondrichthyan species: the Australian ghost shark as a holocephalan reference, and the catshark, devil ray, and little skate as representatives of major elasmobranch lineages. We included the zebrafish (*Danio* rerio) as the ostheochthyan outgroup (Fig. 2C). The ghost shark retains three cone opsin genes—*rh2*, *lws1*, and *lws2*—as well as additional cone-pathway genes such as *gnat2*, *arr3*, *pde6c, cngb3, gngt2* and *grk7*. We therefore assessed a broader panel of cone-associated phototransduction genes across the three elasmobranch genomes, classifying each as retained (Ret), pseudogenized (pseudo), or not recoverable (N.R.) (Fig. 2D). Gene status was assigned on a gene-by-gene basis using a combination of genome annotation, TBLASTN homology searches, and syntenic context where informative. This analysis indicates that the little skate has retained only a limited subset of cone-pathway remnants, with several cone-associated genes represented by pseudogenized loci and others not recoverable as recognizable orthologs in the current assembly.

For *arr3*, *gnat2*, *pde6c*, *cngb3*, and *gngt2*, conserved flanking genes identifiable in ghost shark were also present in corresponding positions in the elasmobranch genomes, permitting synteny-guided comparison. The genomic intervals between these flanking genes were extracted and compared against the ghost shark protein-coding sequences by TBLASTN. For *arr3*, *gnat2*, *pde6c*, and *cngb3*, each little skate locus retained significant similarity to the corresponding cone gene, but the aligned regions were interrupted by nonsense mutations within otherwise conserved coding sequence, consistent with pseudogenization (Fig. S3B, S3C). For *gngt2*, TBLASTN did not recover a significant hit within the conserved syntenic locus and the gene was therefore classified as not recoverable (Fig. S3C). *pde6h* and *guca1c* were not identifiable by keyword search in the ghost shark genome. TBLASTN using human PDE6H and GUCA1C protein sequences did not recover significant matches in the ghost shark genome beyond unrelated background or non-orthologous hits, supporting their classification as not recoverable. *grk7* was annotated as a pseudogene in the little skate genome but was present as an intact gene in the other species examined. By contrast, a synteny-based approach was not possible for the ghost shark cone opsins *rh2*, *lws1*, and *lws2* because consistent flanking annotations were not available across species. Whole-genome TBLASTN searches using ghost shark LWS1, LWS2, and RH2 protein sequences did not recover distinct cone opsin orthologs in the little skate genome (Fig. S3D). The top hits for LWS1 and LWS2 instead corresponded to pinopsin, whereas the top hit for RH2 corresponded to rhodopsin (*rh1*), indicating that separate cone opsin orthologs were not recoverable in the current assembly. Other alignments correspond to characterized unrelated genes (Fig S3D).

These findings indicate that the developing little skate retina retains embryonic expression of the cone-associated regulator *onecut1*, even as rod-associated developmental regulators and rod phototransduction genes increase strongly from embryo to hatchling and remain elevated in the adult. At the same time, comparative genomic analysis shows that many canonical cone phototransduction genes are either pseudogenized or not recoverable in the current little skate genome assembly. This suggests that even if cone genesis pathways that involve *onecut1* are preserved in the skate, functional differentiated cone photoreceptors are not capable of being formed due to disabling of cone specific genes.

### Little skate Onecut1 has two splice isoforms with developmental changes in ratio

To determine whether *onecut1* is expressed as more than one transcript isoform in the little skate retina, we constructed an embryonic retinal cDNA library using oligo-dT primers to isolate mature transcripts from embryonic bulk RNA extracts. Primers specific to untranslated sequences flanking the annotated *onecut1* coding region were used to PCR amplify *onecut1* transcripts. The recovered amplicon was sequenced and identified as skate *onecut1* by NCBI nucleotide BLAST. Relative to the annotated reference sequence in the current little skate genome assembly (Leri_HHJ_1), however, the recovered amplicon contained two unexpected differences (Fig. 3A,B).

**FIGURE 3.**
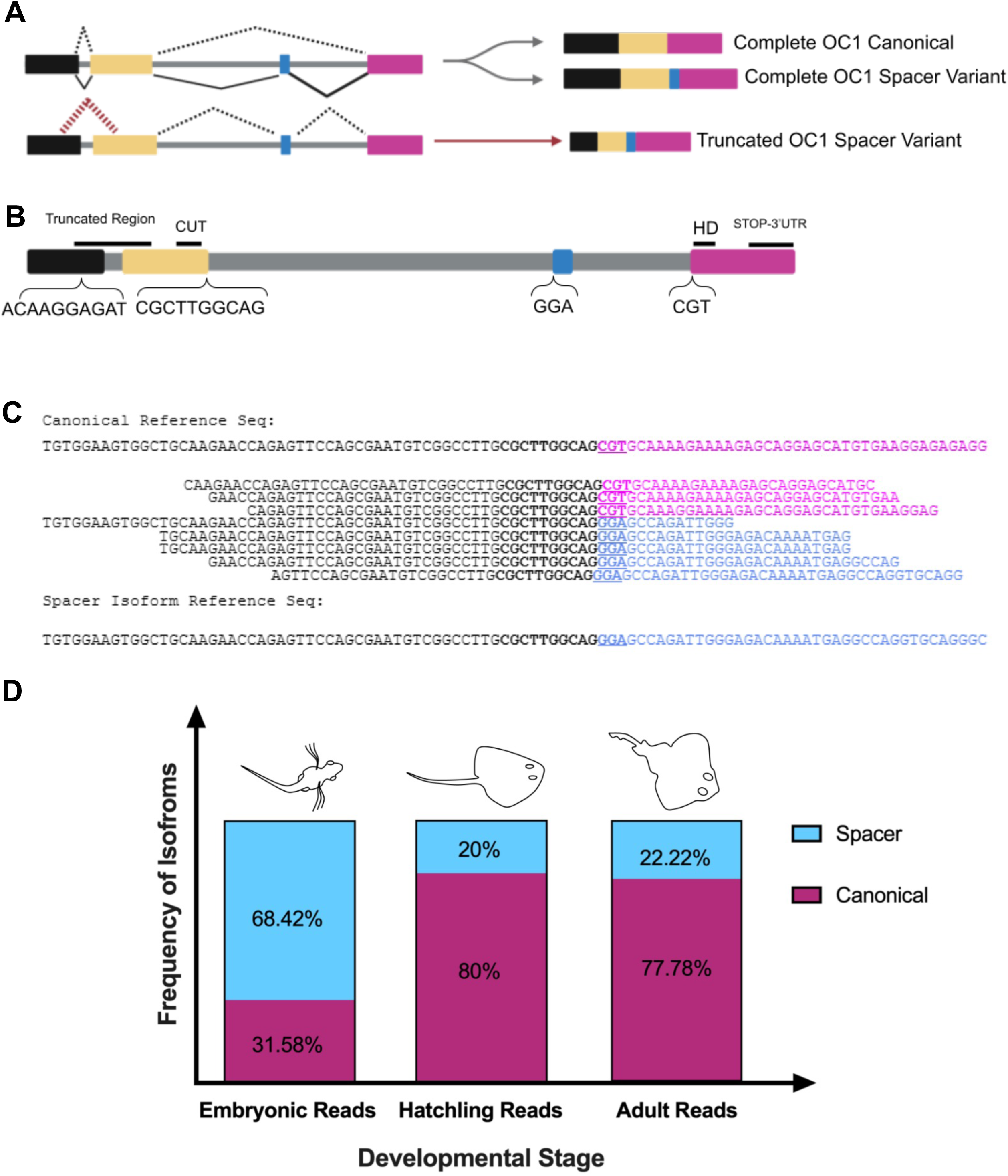
Alternative splicing of Onecut1 results in two isoforms with stage-specific expression ratios. (A) Schematic of the little skate *onecut1* genetic locus. Black box indicates exon 1; yellow box is exon 2; small blue box is the spacer exon; magenta box is exon 3. Observed alternative splicing indicates the presence of two *onecut1* isoforms: canonical *onecut1* and the *onecut1* spacer variant. The third splice event represents the truncated spacer variant that led to the identification of the complete *onecut1* spacer variant. (B) *onecut1* locus with annotated features. Truncation query sequences are labeled in exon 1; spacer query sequence is labeled at the 3’ end of exon 2. Region truncated from canonical *onecut1* spans the first 2 exons (“Truncated Region”). Spacer isoform identifier “GGA” is labeled at the 5’ end of the spacer exon. Canonical isoform identifier “CGT” is labeled at the 5’ end of exon 3. DNA-binding domains--CUT and Homeodomain (HD)--are labeled in their rough relative location within the locus. Distances roughly approximate relative genomic distances between annotated features. (C) (Top) Representative alignment of canonical reference sequence (standard reference’s annotated *onecut1*) with mapped reads containing spacer query (black, bold). (Middle) Magenta sequence shows canonical continuation from the end of exon 2. Bold, underlined “CGT” is 3 bp canonical isoform identifier. Blue sequence shows spacer exon continuing from exon 2, in a majority of reads in the embryo. Bold, underlined “GGA” indicates 3 bp spacer isoform identifier. (Bottom) Alignment of spacer isoform sequence (region of recovered *onecut1* from cDNA library). Blue sequence indicates beginning of spacer exon and bold underlined “GGA” the spacer isoform identifying sequence. (D) Bar plot showing the ratio of isoforms across 3 stages of development. Spacer isoform represented in blue, canonical in magenta. Read counts—Embryo: 75 reads, Hatchling: 25 reads, Adult: 9 reads. Chi-squared = 21.4, 2 df. p < 0.0001

One of these differences was a 144 bp insertion located between the two DNA-binding domains (Fig. 3A,C). The CUT domain of Onecut1 ends at the 3′ end of exon 2, whereas the homeodomain begins at the 5′ end of exon 3. The inserted 144 bp sequence maps to the intron between these exons (Fig. 3B), indicating that it represents a genuine alternative splicing event and defines a previously unannotated exon in the little skate *onecut1* gene. Inclusion of this exon introduces a 48 amino acid spacer between the CUT and homeodomains that is predicted to be intrinsically disordered. A disordered spacer between the DNA-binding domains of Onecut1 has also been described in rat liver^29^, and alternative splicing of linker regions between DNA-binding domains has been reported in other transcription factor families, including *Pou* genes^30^. Notably, BLAST analysis of the recovered skate spacer-containing transcript identified the current *onecut1* annotation of the myliobatiform lesser devil ray (*Mobula hypostoma*) as the closest match, because that annotation likewise predicts a similar spacer-containing coding sequence.

We next asked whether the canonical *onecut1* isoform, lacking the spacer exon, is co-expressed temporally with the spacer-containing isoform in the little skate retina. Using the same RNA-seq read-validation strategy, we confirmed the presence of the spacer element in bulk RNA-seq reads. Reads containing sequence upstream of the insertion site could be assigned either to the canonical splice junction, in which exon 2 splices directly to exon 3, or to the alternative splice junction, in which exon 2 splices to the spacer exon (Fig. 3C). Both splice forms were detected at all three developmental stages examined.

We then characterized the relative abundance of these isoforms across development. Total *onecut1* expression was highest in the S29 embryo (46.09 TPM, compared with 25.28 TPM in hatchling and 6.67 TPM in adult)(Fig. S3E), and the embryonic stage also showed a marked enrichment of the spacer-containing isoform (68.42% spacer across embryonic samples; Fig. 3D, Fig. S4A). By contrast, the spacer isoform was much less abundant in hatchling and adult retina (10% and 11% of *onecut1* transcripts, respectively). Chi-squared analysis indicated that splice isoform usage varies significantly across developmental stages (Fig. 3D).

The second unexpected difference in the recovered amplicon was a 366 bp deletion within the intrinsically disordered region (IDR) of the protein, spanning the first two exons (Fig. 3A,B). This deletion preserved the open reading frame and did not disrupt either of the two DNA-binding domains, the CUT domain or the homeodomain. We therefore asked whether this deletion represented a genuine splice variant or instead arose during library preparation or PCR amplification. To test this, we examined embryonic bulk RNA-seq reads aligning to *onecut1* and isolated reads containing a 10 bp query sequence spanning the putative truncation site. If the truncated isoform were present in vivo, reads containing this query should continue into the downstream sequence recovered from the cDNA amplicon. However, no such alignments were observed: reads containing the query sequence did not align with the truncated region of the recovered isoform (Fig. S4B). These findings suggest that the recovered IDR truncation is most likely an artifact introduced during cDNA synthesis or downstream amplification.

Together, these findings identify a previously unannotated, developmentally regulated spacer exon in little skate *onecut1*. Whereas the recovered IDR-truncated amplicon is most consistent with a library or amplification artifact, the 144 bp spacer insertion is supported by RNA-seq reads and is present alongside the canonical splice form at all stages examined. The spacer-containing isoform is strongly enriched in the embryonic retina, when overall *onecut1* expression is also highest, suggesting that alternative splicing of *onecut1* is developmentally regulated in the little skate. These results motivated functional testing of the canonical and spacer-containing isoforms in subsequent experiments.

### LSOC1X1 and LSOC1X2 Activate ThrbCRM1 in the P0 Mouse

Because the canonical developmental role of Onecut1 in other vertebrates includes activation of *thrb* and other cone-associated genes^16,31^, the developmental enrichment of the spacer-containing isoform raised the question of whether inclusion of a 48 amino acid spacer between the DNA-binding domains might alter Onecut1 activity. Because the cDNA library recovered only the IDR-truncated product, we synthesized full-length gene fragments corresponding to canonical little skate Onecut1 (LSOC1X1) and spacer-containing little skate Onecut1 (LSOC1X2), which differs from the canonical transcript only by inclusion of the 144 bp spacer exon. These gene fragments were cloned into expression vectors for subsequent functional analysis.

To test whether the skate Onecut1 isoforms retained the ability to activate a cone-associated reporter, we ex vivo electroporated expression constructs for the canonical isoform (LSOC1X1) and the spacer-containing isoform (LSOC1X2) into P0 mouse retinas together with the ThrbCRM1::GFP reporter. ThrbCRM1 has two possible Otx2 binding sites and one Onecut1 binding site, and this binding-site organization is conserved across several terrestrial vertebrates, from mammals to frogs (Fig. 4A)^16^. In this assay, exogenous mouse Onecut1 robustly activates ThrbCRM1, whereas electroporated cells in the absence of exogenous Onecut1 do not, because the retinal cells targeted at this stage do not normally express *Onecut1* and therefore do not activate the reporter. Reporter activation in this paradigm is also dependent on Otx2, indicating that the assay reveals the cooperative activity of Onecut1 and endogenous mouse Otx2^16^.

**FIGURE 4.**
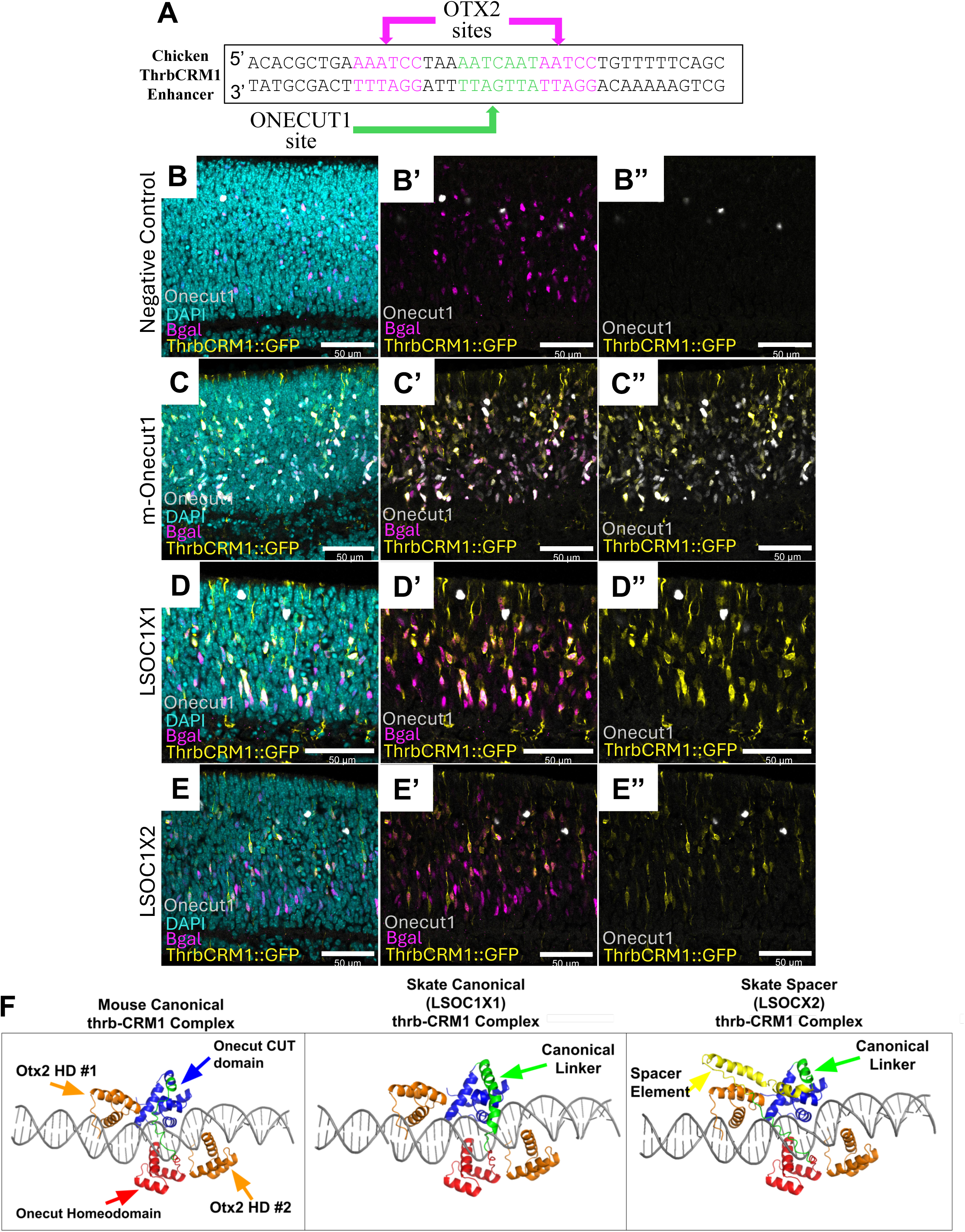
Novel Onecut1 spacer isoform validated as functional protein, activates the thrbCRM1 reporter element. (A) Annotated ThrbCRM1 enhancer sequence, adapted from Emerson et al. (2013), showing the predicted Onecut1 and Otx2 binding sites. The Onecut1 binding site is marked with a green arrow, while the two Otx2 binding sites are marked with purple arrows. (B-E) P0 mouse retinas electroporated ex vivo with ThrbCRM1::GFP, pCAG::Bgal, and a pCAG construct driving expression of the Onecut1 transcript denoted at the left of each row. After 2 days, retinas were harvested and processed for confocal microscopy to visualize nuclei (DAPI in blue), GFP reporter (yellow), Bgal (magenta), and Onecut1 protein (white), which is detected in unelectroporated mouse horizontal cells in all retinas and in any electroporated cell expressing a Onecut1 protein recognized by the antibody. Visualized channels are shown above each row. (B) pCAG::NULL negative control: pCAG expression vector with no encoded protein sequence. (C) pCAG::mouse Onecut1. (D) pCAG::LSOC1X1. (E) pCAG::LSOC1X2. Scale bars are all 50 μm. (F) AlphaFold3 models predicting the binding properties of one variable Onecut1 molecule (mouse, skate canonical, or skate spacer isoform), two molecules of Otx2, and the ThrbCRM1 double helix. All DNA-binding domains, the Onecut1 linker, spacer, and ThrbCRM1 are shown. Onecut1 CUT (blue), HD (red), spacer element (yellow), and canonical linker (green) are labeled. Otx2 HD is labeled in orange.

The skate isoforms were cloned into a CAG-based misexpression vector with broad activity in dividing retinal cells and their progeny ^32^. P0 mouse retinas were electroporated in four groups: CAG::mouse Onecut1, CAG::LSOC1X1, CAG::LSOC1X2, and negative control CAG::NULL. CAG::nuclear β-galactosidase (β-gal) was co-electroporated as an electroporation marker, together with ThrbCRM1::GFP as the reporter of Onecut1/Otx2-dependent activity. After two days in culture, retinas were processed for confocal microscopy to visualize electroporated cells (β-gal), reporter activation (GFP), and Onecut1 immunoreactivity.

As expected, negative-control retinas showed no induced ThrbCRM1::GFP signal and no ectopic Onecut1 immunoreactivity beyond that detectable in endogenous horizontal cells (Fig. 4B). By contrast, misexpression of mouse Onecut1 robustly induced ThrbCRM1::GFP and produced strong Onecut1 immunoreactivity throughout the electroporated region (Fig. 4C). Importantly, both LSOC1X1 and LSOC1X2 also induced ThrbCRM1::GFP expression (Fig. 4D,E). Neither skate isoform was detected by the mouse Onecut1 antibody, consistent with the lack of cross-reactivity of this reagent to the skate protein (Fig. 4D”,E”). These results show that both skate Onecut1 isoforms retain the ability to activate the ThrbCRM1 reporter in a heterologous mouse retinal context, indicating that the spacer-containing isoform is not functionally inactivated with respect to this assay.

To provide a structural framework for interpreting this result, we then modeled complexes containing ThrbCRM1, two molecules of mouse Otx2, and either mouse Onecut1, LSOC1X1, or LSOC1X2 using AlphaFold3, which has recently been shown to predict multi-molecular interactions with high accuracy^33^. Our modeling assumptions were intentionally minimal and were based on prior experimental evidence that Onecut1 and Otx2 cooperate to activate ThrbCRM1^16,23^. We therefore modeled a reduced complex containing the ThrbCRM1 DNA element, one Onecut1 molecule, and two Otx2 molecules, reasoning that this was the simplest biologically motivated configuration in which to ask whether inclusion of the skate spacer exon would favor a qualitatively altered binding arrangement. The goal of this analysis was not to reconstruct the full in vivo regulatory complex, but to test whether spacer inclusion alone was sufficient to produce an obvious structural disruption under a minimal cooperative binding scenario. In the control model containing mouse Onecut1, the DNA-binding domains were positioned in contact with their expected binding regions on the ThrbCRM1 element (Fig. 4F, left). Substitution of mouse Onecut1 with LSOC1X1 or LSOC1X2 did not produce a qualitatively different global binding arrangement (Fig. 4F, middle and right). Although the spacer insertion in LSOC1X2 introduces additional predicted disorder between the CUT and homeodomains, the model did not suggest a major steric disruption of the overall complex under the conditions tested. Because the LSOC1X2 model did not adopt a qualitatively different global arrangement from the canonical model, we interpret the spacer as unlikely to abolish ThrbCRM1-associated activity by forcing a different binding mode.

Together, these findings show that both the canonical and spacer-containing little skate Onecut1 isoforms retain the ability to activate ThrbCRM1 in a heterologous mouse retinal assay. This simply demonstrates the ability of these two isoforms to activate transcription. The inability of the mouse monoclonal antibody to detect LSOC1X1 or LSOC1X2 confirms that this reagent is not suitable for assessing skate Onecut1 protein expression but does not affect the reporter-based functional readout. AlphaFold3 modeling further suggests that inclusion of the spacer exon does not impose a qualitatively different global binding arrangement at the ThrbCRM1 element. Thus, under the conditions tested, the spacer-containing isoform remains compatible with ThrbCRM1-associated activity.

## Discussion

The little skate retina provides a relatively unique opportunity to examine how a vertebrate visual system can lose canonical cone photoreceptors while retaining aspects of photopic function. In most studied vertebrates, the mature retina contains rods together with one or more cone subtypes, and photoreceptor development proceeds through partially overlapping but ultimately divergent gene regulatory programs^24^. By contrast, the little skate appears to mature with only rods, yet those rods display light-adaptive properties that extend beyond the typical vertebrate rod response profile^10^. Our results show that the little skate retina is experimentally tractable during development, and they identify a molecular context in which cone-associated regulatory features are retained even as the mature retina adopts a rod-only identity.

A first contribution of this study is methodological. We show that the embryonic little skate retina can be accessed experimentally through EdU birthdating, immunohistochemical analysis, and retinal electroporation. These approaches allowed us to identify S29 as a stage containing actively dividing retinal cells and Otx2-positive cells consistent with early photoreceptor production. The successful electroporation of embryonic skate retina establishes a platform for future functional studies in a species for which developmental gene regulation remains poorly characterized. At the same time, the lack of ThrbCRM1 reporter activation in the skate retina under the conditions tested should be interpreted cautiously. This result may reflect absence or alteration of the Otx2/Onecut1 regulatory state described in other vertebrates, but it could also arise from species-specific differences in enhancer interpretation, developmental timing, chromatin state, or required cofactors that should be investigated in future studies.

Our expression and comparative genomic analyses indicate that the little skate retina retains some early cone-associated regulatory features while showing extensive loss of canonical cone-pathway components. *onecut1* is most highly expressed in the embryo and declines thereafter, whereas rod-associated developmental regulators, including *mafb* and *nr2e3*, increase from embryo to hatchling and remained elevated in the adult. Rod phototransduction genes show the same general trajectory, increasing strongly as development proceeds toward hatching. In contrast, comparative genomic analysis across chondrichthyans indicate that many canonical cone phototransduction genes in the little skate are either pseudogenized or are not recoverable as recognizable orthologs in the current assembly. Together, these findings argue against a simple model in which the cone developmental program was entirely erased. Instead, they suggest a more selective rewiring, in which some upstream or early regulatory features persist while much of the canonical cone differentiation machinery has been lost or degraded.

The developmental behavior of *onecut1* is especially notable in this context. In other vertebrates, Onecut1 has a well-established role in cone, horizontal and ganglion cell development and participates in regulatory interactions that oppose rod specification ^13–16,34^. Perturbation studies in mouse and chick have shown that Onecut1 can promote cone-associated programs and suppress rod-associated ones, while rod-specifying factors such as Nrl can redirect developing photoreceptors toward rod fate ^16,34,35^. In the little skate, however, the embryonic enrichment of *onecut1* occurs in a retina that ultimately lacks cones. Bulk RNA-seq cannot determine which cells express *onecut1*, nor whether Onecut1 and Otx2 are co-expressed in the same embryonic progenitors. Accordingly, our results do not show that Onecut1 directly drives a cone-like developmental state in skate retinal progenitor cells. They do show, however, that *onecut1* expression is retained in the embryonic retina and is developmentally regulated in a way that makes it a plausible component of the altered photoreceptor regulatory network in this lineage.

Our analysis of large Maf family members identified three large Maf genes in the little skate: *mafaa, mafb*, and *c-maf.* By contrast, we did not recover a distinct *nrl* gene ortholog from the current little skate genome assembly. Because *nrl* is widely treated as a central regulator of rod specification in mammals, its apparent absence in the skate is notable. Among the large Maf family members that were identified, *mafb* showed the highest expression across developmental stages, making it the strongest candidate for large-Maf-associated involvement in skate rod development. These findings establish that the skate rod developmental program is deployed in the absence of a recognizable *nrl* gene ortholog and provide a basis for further comparison with large Maf evolution in other vertebrate lineages. It was previously reported that the four large Maf orthology classes were established before the divergence of actinopterygians and sarcopterygians, and further suggested that these classes may have arisen prior to the gnathostome radiation^26^. Under that interpretation, the absence of a distinct nrl ortholog in the little skate would most likely reflect lineage-specific loss. However, the targeted genome analyses used in the present study were designed to assess gene presence, absence, and pseudogenization within a limited comparative set, and are not sufficient to resolve the deeper evolutionary history of the *Nrl* orthology class. Determining when *Nrl* arose, and whether it was ancestrally present in chondrichthyans more broadly, will require broader phylogenetic sampling, including cyclostomes and a wider range of elasmobranch genomes, as noted by Coolen et al^26^.

A central finding of this study is that little skate *onecut1* undergoes developmentally regulated alternative splicing to produce a previously unannotated spacer-containing isoform. The presence of a spacer-containing Onecut1 isoform is intriguing because similar linker-modifying splicing has been reported in other transcription factor families, and because Onecut1 itself is known to be alternatively spliced in rat liver ^29,30^. In those contexts, isoform-specific differences in promoter responsiveness suggest that alternative splicing can alter DNA-binding preferences or regulatory selectivity without abolishing activity^29^. Transcript reads of *Onecut1* sourced from two publicly available mouse retinal datasets show low level splicing in a region corresponding to rat *Onecut1* alternative exon identified in previous study, which also introduces a spacer in between the DNA binding domains (Fig S4C-E). It is unclear if any of these correspond to a complete alternative transcript, or if the level to which it is expressed has a meaningful impact on regulation. Several myliobatiformes have an annotated *onecut1* isoform containing the 48 amino acid spacer motif present in the skate (Fig S3F). Tissue specific expression studies in this clade are lacking and would allow us to determine if alternative splicing of *onecut1* in the manner observed is a broader batoid phenomenon. Inspection of Onecut1 transcripts from shark retinal transcriptomes, such as S. canicula (recently analyzed by single-nucleus sequencing), may also be informative, although the current reference genome assembly for this species does not predict the observed spacer splicing pattern^36^.

Our explant assay in the P0 mouse retina indicates that both the canonical skate isoform (LSOC1X1) and the spacer-containing isoform (LSOC1X2) retain the ability to activate ThrbCRM1 in a heterologous retinal context. This result is important because it argues against the simplest loss-of-function model: inclusion of the 48 amino acid spacer does not render the protein incapable of enhancer-associated activity in this assay. At the same time, the assay should not be interpreted as a direct readout of endogenous skate gene regulation. It instead shows that both skate proteins remain functionally competent to cooperate with the mouse retinal environment, including endogenous mouse Otx2, to activate a known Onecut1-responsive reporter.

The AlphaFold3 models are best interpreted in the same limited but useful framework. Our modeling assumptions were intentionally minimal and were based on prior experimental evidence that Onecut1 and Otx2 cooperate to activate ThrbCRM1^16^. We therefore modeled a reduced complex containing the ThrbCRM1 DNA element, one Onecut1 molecule, and two Otx2 molecules, not to reconstruct the full in vivo regulatory complex, but to ask whether inclusion of the spacer exon alone would favor a qualitatively altered global binding arrangement, which it did not. The spacer-containing model also did not adopt a grossly different overall configuration from the canonical model, consistent with the explant result showing preserved reporter activation. These models do not establish the true stoichiometry, dynamics, or mechanism of the endogenous skate complex, but they do support the narrower inference that spacer inclusion is unlikely to abolish ThrbCRM1-associated activity by forcing an obviously incompatible binding mode.

Taken together, these findings suggest that the loss of cones in the skate lineage is unlikely to be explained by a simple absence of *onecut1* coding potential or by complete inactivation of Onecut1 protein function. A broader regulatory explanation is more likely. One possibility is that *onecut1* is expressed in an altered cellular context relative to other vertebrates or other necessary cone genesis components are disabled. Another is that the downstream cis-regulatory architecture of cone-associated genes, including but not limited to *thrb*, has diverged such that the same upstream factors no longer drive the same targets. A third possibility is that skate photoreceptor development involves a partial uncoupling of early cone-associated developmental features from terminal cone differentiation, allowing certain cone-like features to be deployed in cells that ultimately mature as rods. Our data do not distinguish among these possibilities, but they narrow the problem by identifying a specific developmental regulator whose transcript structure and developmental deployment differ from those in the best-studied vertebrate systems.

An attractive speculative model is that early skate photoreceptor precursors transiently engage parts of a cone-associated regulatory program, potentially including *onecut1*, while later developmental events channel those cells toward rod identity and rod outer segment maturation. Under such a model, the spacer isoform might modify the transcriptional selectivity of Onecut1 rather than simply eliminating its activity, perhaps allowing some developmental or synaptic features associated with cone-like programs to be retained without full commitment to cone identity. This idea remains hypothetical, and we have not shown that the spacer isoform acts in that way in vivo. Nonetheless, it offers one possible explanation for how the skate retina could combine rod morphology and rod phototransduction with functional and structural features that appear, in some respects, cone-like.

More generally, the little skate retina illustrates how evolution may reshape cell identity not only through gene loss, but also through altered deployment of conserved developmental regulators and modulation of transcript structure. In that sense, the skate is not merely a cone-less oddity. It is a system in which the relationship between photoreceptor identity, enhancer logic, and developmental gene regulation can be examined in an evolutionary context. Our results identify a developmentally regulated *onecut1* splice isoform, establish functional tools for embryonic skate retina, and define a comparative molecular framework for studying how a rod-only vertebrate retina was built.

## Methods

### Preparation of Ringer’s Solution

All skate Ringer’s solutions were prepared as follows: 10 g Urea, 7 g NaCl, and 2 g trimethylamine N-oxide (TMAO) were dissolved in 500 ml of RO water, and filter sterilized. “(Ringer’s Solution)” indicates that the solution referred to was supplemented with this same concentration of solutes, dissolved in the standard solution. Where this label is not given, the solutions were prepared standard.

### Tissue Preparation

Retinas were dissected from embryonic specimens in 1XPBS (Ringer’s Solution), ensuring the preservation of the vitreous layer. The tissues were immediately fixed in 4% paraformaldehyde in 1X PBS (Ringer’s Solution) for 30 minutes. The tissue was washed with PBS (Ringer’s Solution) 3 times and submerged in 30% sucrose in 1x PBS (Ringer’s Solution) and stored at 4°C overnight to allow for complete equilibration. The following day, sunken retinas were removed from the sucrose solution, embedded in optimal cutting temperature (OCT) compound, and rapidly frozen at -80°C. Cryosections of 20 µm thickness were obtained from the frozen tissue using a Leica cryostat. The sections were air-dried for 20 minutes and then stored at -80°C until further processing.

### Antibody Staining

Frozen tissue sections were thawed in 1XPBS at room temperature prior to staining. Blocking was performed by incubating the sections in 5% serum (either goat or donkey, depending on the origin of the secondary antibody) diluted in PBT (0.1% Tween 20 for skate samples or 0.3% Triton-X in PBS for mouse tissue) for 1 hour. The sections were then incubated overnight at 4°C in the dark with primary antibodies diluted in 5% blocking solution (Normal Goat Serum) in PBT. The following day, the slides were washed three times for 15 minutes each with PBT. A second blocking step was performed by incubating the sections for 30 minutes in a 5% blocking solution in PBT. The slides were then washed three times for 5 minutes each with PBT and incubated overnight at 4°C in the dark with secondary antibodies diluted in PBT (without serum). On the third day, slides were washed again with PBT (two washes of 15 minutes each) and a final wash with PBT containing 0.1% DAPI, protected from light. After removing the solution, the slides were mounted in Fluormount-G (Southern Biotech), cover-slipped, sealed with nail polish, and allowed to dry for 15 minutes to 1 hour in a fume hood. The slides were stored at 4°C before imaging.

#### Antibodies

**Antibodies:**
| **AB** | **Type** | **Catalog number** | **Vendor** |
| --- | --- | --- | --- |
| **Otx2** | Goat | AF1979 | R&D Systems |
| **Onecut1** | Mouse | sc-376167 | Santa Cruz Biotechnology |
| **Beta Gal** | Mouse | DSHB-40-1a | Developmental Studies Hybridoma Bank |
| **Beta Gal** | Chicken | ab9361 | Abcam |
| **GFP** | Chicken | ab13970 | Abcam |
| **AU1** | Mouse | 901901 | Biolegend |

### Microscopy Imaging

Images were collected using a Zeiss LSM 880 confocal microscope. Fluorescence channels employed included Alexa Fluor 647, Alexa Fluor 488, Cy3, and DAPI, with the specific channel used indicated in the corresponding Results sections. Laser intensity was adjusted on a case-by-case basis to maximize signal-to-noise ratio, ensuring that the signal was clearly distinguishable from background fluorescence, with care to limit bleed-through effects. Post-processing of images was performed using ImageJ (NIH). Adjustments included brightness and contrast optimization, and color changing to enhance image clarity and ensure consistency of color scheme. Adjustments were applied equally across an image.

### Embryo Selection

Embryonic little skates (*Leucoraja erinacea*) were selected for analysis based on their developmental stage, determined by visual inspection (candling). Embryos were staged according to images and described landmarks in little skates and winter skates^17,18^. At the S29 stage, the embryos exhibit distinctive features such as exterior gill filaments and protruding limb buds. It should be noted that these time points are estimates and may vary slightly between individuals.

### Euthanasia

To euthanize the selected embryos, they were submerged in a solution containing 1g/L Syncaine in seawater, supplemented with 2g/L sodium bicarbonate. The embryos were kept in this solution until cessation of movement and confirmed to be non-responsive and decapitated with scissors.

### EdU injection

For EdU incorporation studies, 20 µL of 10 mM EdU was injected into the yolk of embryonic (∼50 days post-oviposition) or hatchling little skate (*Leucoraja erinacea*). The injections were targeted to the remaining yolk on the underside of the hatchlings, ensuring proper exposure to the EdU. Following injection, the skates were allowed to recover in a seawater table for selected lengths of time before sacrifice and dissection as per the experimental design. EdU+ cells were detected using the Click-iT EdU Alexa Fluor 647 imaging kit (Invitrogen, Carlsbad, CA, USA), following the manufacturer’s protocol. This method allowed for precise identification of cells that had incorporated EdU during DNA synthesis.

### Anesthesia for EdU Yolk Injection

For EdU yolk injections, embryos were anesthetized by transfer to a solution of 10 mg Syncaine and 20 mg sodium bicarbonate dissolved in 100 ml of seawater. The embryos were monitored for a decrease in movement to ensure effective anesthesia before proceeding with injections.

### EdU Labeling Detection

For EdU labeling detection, the protocol was adapted from the immunohistochemistry (IHC) procedure previously described, as EdU detection was performed alongside antibody staining. On the third day of the IHC protocol, after the initial 2x PBT washes, the slides were washed once with 1XPBS. A detection solution was then prepared according to the manufacturer’s direction. This solution was applied to each slide at 200 µL per slide, followed by a 30 minute incubation. After incubation, the solution was removed, and the slides were washed 2X with PBT for 15 minutes each, followed by a final wash with PBT containing 0.1% DAPI. Mounting as above under “Antibody Staining”.

### RNA Isolation and Quality Control

RNA was isolated with a Promega Maxwell RSC miRNA tissue kit. The RNA quality of the bulk samples was assessed using an Agilent Bioanalyzer, RIN values ranged from 8.3 to 10, indicating high RNA quality across all samples. The concentrations, as measured by the Bioanalyzer, varied from 24.4 ng/µL to 67.9 ng/µL, with corresponding total RNA amounts ranging from 610 ng to 1697.5 ng.

### Paired-End Sequencing

RNA samples were sequenced on the Illumina platform (Columbia Genome Center). Paired-end sequencing was performed, generating reads with lengths 10-75 base pairs. RNA-Seq data can be found at NCBI GEO as GSEXXXXX, available after publication.

### Sequencing Quality Control

Brief quality control metrics for each sequencing run are as follows:

- Embryonic Samples (CM003-CM005):

◦ Total Sequences: 20.99-24.60 million per run
◦ GC Content: 45-46%
- Hatchling Samples (CM006-CM008):

◦ Total Sequences: 19.79-22.68 million per run
◦ GC Content: 44-45%
- Adult Samples (CM009-CM010):

◦ Total Sequences: 24.49-29.88 million per run
◦ GC Content: 45%

All reads were analyzed for quality, and no sequences were flagged as poor quality across all samples.

### Data Processing

The reads were aligned to the genome listed under the “Annotation” section, using the STAR aligner with two-pass mode (basic), with specific splice junction detection parameters set according to the quality of the reads and the reference genome. The resulting BAM files were then sorted and indexed for downstream analysis.

### Annotation

The gene annotation file used for this project was based on the Leri_hhj_1 genome assembly (NCBI Assembly Accession: GCF_028641065.1). This annotation, which follows the GTF version 2.2 format, was provided by NCBI RefSeq with the genome build annotated on April 21, 2023. The genome assembly and the corresponding gene annotation were both sourced using the NCBI Genome Data Viewer UI. The annotation source is listed as “NCBI RefSeq GCF_028641065.1-RS_2023_04.” As many genes are listed by location number in the annotation file, a combined gene matrix was created consisting of location labeled genes (i.e. those that appear as LOC#########) annotated with a long form alias, sourced in batch using NCBI’s Entrez gene feature. As no short form alias was located for these genes, they were not imported into the large objects used in bioinformatic processing, but the matrix was used to identify relevant genes (r*hodopsin, mafb*). Table available on request. Gene names throughout the text are lowercase, to reflect the conventions of the datasets used to perform analysis on the skate.

### Gene Nomenclature

Gene symbols are formatted according to species-specific annotation conventions and are italicized throughout the manuscript. For little skate and other non-mammalian vertebrates analyzed here, including zebrafish, catshark, devil ray, and ghost shark, gene names are given in lowercase italics (for example, *onecut1, mafb,* and *thrb*), reflecting the conventions used in the underlying genome annotations and transcriptomic datasets. Mouse gene symbols are written with an initial capital letter and the remaining letters lowercase, also in italics (for example, *Onecut1, Mafb,* and *Thrb*). Human gene symbols, where referenced, are written in uppercase italics.

Protein names are not italicized. Thus, proteins are referred to in roman type, using conventional capitalization appropriate to the context (for example, Onecut1, Otx2, and NRL). Reporter constructs, enhancers, and plasmid names are also written in roman type (for example, **ThrbCRM1**, **ThrbCRM1::GFP**, and **CAG::mCherry**). This convention is used throughout the manuscript to distinguish genes, proteins, and experimental constructs clearly.

#### Data Analysis for Bulk RNA Sequencing

For bulk RNA sequencing analysis, gene counts were generated for each sample using the FeatureCounts tool available on the Galaxy bioinformatics platform. This process was repeated for each sequencing run. Following the feature count extraction, the counts from the two sequencing runs for each sample were averaged. The resulting averaged counts were then converted into Transcripts Per Million (TPM) to normalize the gene expression levels across samples. TPM normalization was performed by calculating the read counts per kilobase of transcript per million mapped reads for each gene, and then scaling these values such that the sum of all TPM values in a sample equaled 1 million. After calculating TPM for each sample, the average TPM and standard error were computed within each experimental group (e.g., embryo, hatchling, and adult). Gene expression across developmental stages was visualized using two heatmaps generated in R with the pheatmap package. For each gene of interest, average TPM values were extracted for each developmental stage (embryo, hatchling, adult) and log2-transformed with a pseudocount of 1 [log2(TPM + 1)] to compress the dynamic range while preserving absolute expression levels. Color intensity reflects absolute expression magnitude—color matching the number n on the scale bar is expressed at an average of 2^n TPM across the samples for the stage group. A diverging color scale from steelblue (low) through light yellow to firebrick (high) was used, spanning 0–8 log2 TPM for the developmental transcription factor heatmap (Fig 2A) and 0–12 log2 TPM for the phototransduction gene heatmap (Fig 2B). Genes were hierarchically clustered using euclidean distance and complete linkage. They were annotated by pathway category (rod, cone, or shared) (add lamb citation). Previously reported mouse datasets visualized for the Onecut1 genomic region were GSM2720093 (E14.1; Mus musculus; RNA-Seq; SRR5877176) and GSM2319611 (E14.5 replicate 1; Mus musculus; RNA-Seq; SRR4253086)^37,38^.

#### cDNA Library Construction

First-strand cDNA synthesis from the RNA samples used for the bulk RNA-seq processing was initiated by mixing 7 µL of RNA with 2 µL of a primer mix (Invitrogen, Catalog #18418012, Lot #1793155) and 1 µL of 100 mM dNTP mix (Agilent, Lot #200418-51), making up a total volume of 10 µL. The mixture was incubated at 65°C for 5 minutes, then rapidly chilled on ice. Subsequently, 4 µL of 5x Protoscript II buffer (NEB, Catalog #B0368S, Lot #10118640), 2 µL of 0.1M DTT (NEB, Catalog #B1043A, Lot #10122162), 1 µL of Protoscript II reverse transcriptase (20 U/µL) (NEB, Catalog #M0368S, Lot #10125099), 0.2 µL of RNase inhibitor (Invitrogen, Catalog #10777-019, Lot #1840422), and 2.8 µL of nuclease-free water were added to the RNA-primer mix, making the final volume 20 µL. The reaction was incubated at 42°C for 1 hour, followed by a final incubation at 65°C for 20 minutes to terminate the reaction.

### Molecular Biology

To generate a little skate misexpression construct, we isolated Onecut1 cDNA from both the embryonic and adult stages of the little skate retina. Total RNA was extracted, reverse-transcribed into cDNA, and the gene was PCR-amplified. The skate Onecut1 sequence was amplified from the cDNA library using specific primers designed to flank the coding region. The forward primer used was **GATCTCACAGAGAAAGGAC**, and the reverse primer was **ACTAGTGATCACCGAGG**. PCR amplification was carried out using Herculase polymerase with an initial denaturation step at 95°C for 3 minutes, followed by 35 cycles of 95°C for 30 seconds, 58°C for 30 seconds, and 68°C for 75 seconds, with a final extension at 68°C for 6 minutes. The resulting PCR product was then purified and used for subsequent cloning and expression studies. The resulting PCR product was then A-tailed through incubation with Taq polymerase and dATP and cloned into the pGEM-T Easy vector. The Onecut1 insert was cut out using EcoR1 was then ligated downstream of the CAG element in a misexpression vector. For experiments involving synthetic gene fragments, full-length skate onecut1 canonical (LSOC1X1) and spacer (LSOC1X2) isoforms were obtained from Twist Bioscience, each designed with EcoRI and NotI overhangs at the 5′ and 3′ ends, respectively. Fragments were amplified using universal adapter primers and ligated into pCAG::GFP using EcoR1/Not1 and replacing GFP. Clones were confirmed by sequencing and DNA for electroporation was produced using a Qiagen Plasmid Midi Kit.

### Electroporations

For both mouse and skate, electroporations followed the general procedure as used previously^39^ with the following modifications and details. <u>Skate</u>: Retinas were harvested in Ringers L15 media and electroporation mixes were made with 40 µl 0.96M D-mannitol, 7.5 µl 10X PBS, and 10 µg each plasmid diluted in a total of 10 µl Tris-EDTA (TE) buffer. After electroporation, retinas recovered in Ringers L15 media supplemented with 10% Fetal Bovine Serum, L-glutamine, penicillin, and streptomycin and then were placed on 13mm/0.2 micron filters (10417001, Cytiva) floating on the same supplemented media. Culture was carried out at ∼22 degrees for three days. <u>Mouse</u>: Retinas were harvested in DMEM/F12 and 5 µg of each plasmid was combined in a total volume of 50 µl 1XPBS for electroporation. A total of four retinas were electroporated for each condition and cultured at 37 degrees for two days before harvest. All electroporations used a Nepagene NEPA21 Type II Super Electroporator with 25V pulses, 50ms pulse interval, 950ms interpulse interval, for 5 total pulses.

### AlphaFold Structural Modeling

AlphaFold3 models were generated using the Google AlphaFold server. We modeled:

1. Skate canonical Onecut1
2. Skate spacer Onecut1
3. Mouse canonical Onecut1
4. Truncated library-derived skate Onecut1

Sequences were sourced from NCBI Genome Data Viewer. Models included two mouse Otx2 protein sequences and the chick thrb-CRM1 sequence (40 bp upstream regulatory element: ACACGCTGAAAATCСТААААТСААТААТCCTGTTTTTCAGC). The reverse complement was also provided to model DNA as a double helix. Complexes were visualized in PyMOL to examine DNA binding domains.

### TBLASTN Analysis

Phototransduction genes were identified across four chondrichthyan genomes—little skate (Leucoraja erinacea, GCF_028641065.1), small-spotted catshark (Scyliorhinus canicula, GCF_902713615.1), Atlantic devil ray (Mobula hypostoma, GCF_963921235.1), and ghost shark (Callorhinchus milii, GCF_018977255.1)—by first searching the NCBI Genome Data Viewer for annotated genes and pseudogenes using gene names and descriptive keywords, followed by whole-genome TBLASTN with ghost shark orthologous proteins for genes not recovered by annotation (NCBI BLAST+ v2.17.0; E-value threshold 1×10⁻³). When a ghost shark ortholog was not clearly identifiable, the corresponding human protein sequence was obtained from the NCBI RefSeq database for the human reference assembly GRCh38.p14 and used in a TBLASTN search against the ghost shark genome to determine whether a detectable chondrichthyan sequence could first be established in the reference species. Hits were ranked by bit score, inspected in Genome Data Viewer, and evaluated against the expected syntenic neighborhood defined by flanking ghost shark genes. To confirm hits in the predicted locus, local TBLASTN was performed on extracted intervals spanning the locus. Fragmented multi-exon hits containing in-frame stop codons were interpreted as pseudogene remnants, whereas high-scoring hits outside the expected syntenic region, including hits to paralogs or unrelated genes, were not considered evidence of locus recovery. To identify nrl-related large MAF-family orthologues, additional whole-genome TBLASTN searches were performed using zebrafish Nrl and human NRL protein sequences, and resulting hits were interpreted by ranking alignment scores and inspecting the annotated loci to distinguish mafaa, mafb, c-maf, and related family members.

### Silhouette Images

Fig 2 silhouettes were all taken from Phylopic.org: “*Danio rerio* by Ian Quigley, via PhyloPic.” Licensed under CC BY 3.0.”, “*Callorhinchus milii* by Ingo Braasch, via PhyloPic. Dedicated to the Public Domain (CC0 1.0).”, “*Scyliorhinus canicula* by Birgit Lang, via PhyloPic. Dedicated to the Public Domain (CC0 1.0).”, “*Mobula mobular* by David Orr, via PhyloPic. Public Domain Mark 1.0. The little skate image was hand drawn with google draw.

## Acknowledgements

Funding for this project was provided to M.E. by NSF CAREER Award 1453044, PSC-CUNY award ENHC-54-105, and the MBL Whitman Fellows Program with support from the Hartline MacNichol Research Award and the Fries Trust Research Awards, L.&A Colwin Summer Research Fellowship Fund. D.M. was supported by a Graduate Research Training Initiative for Student Enhancement grant from the National Institute of General Medical Sciences to The City College of New York (5T32GM136499-05). In addition, funding was provided in part through an NIH/NCI Cancer Center Support Grant P30CA013696. We thank Laura Alexander for assistance with RNA isolation, Miruna Ghinia-Tegla for technical guidance on bulk-RNA sequencing, Angelina Grebe for consultation on the potential implications of this discovery at the structural level, Michael Palmer for skate tissue culture assistance, Ken Hastings for marine electroporation guidance, and Sruti Patoori for preliminary investigation of the skate retina. We thank the MBL staff of the Marine Resource Center including Nolan Gibbons, Brian McGonagle, Lisa Abbo, and Dave Remsen for assistance with the use and care of specimens. We thank all members of the Emerson Lab for support throughout the project.

## SUPPLEMENTAL FIGURE LEGENDS

**<u>SUPPLEMENTAL FIGURE S1 Extended EdU/PHH3 time-course analysis and electroporation controls in the embryonic little skate retina.</u>**

(A) Table showing counts of PHH3+ cells and EdU+ cells in the 4-hour and 8-hour post-injection groups. (B) Representative images from the 2-hour and 40-hour EdU groups. Arrowheads mark PHH3+ cells. Top left: 2-hour group showing PHH3+ (white) cells and EdU (magenta); top right: same field with PHH3 channel off showing absence of EdU signal at arrowhead positions, indicating no EdU incorporation in PHH3+ cells at this early time point. Bottom left: 40-hour group with PHH3 staining (magenta) and EdU (white); bottom right: same field with PHH3 channel turned off and EdU signal (white) visible, demonstrating that all PHH3+ cells are also EdU+. (C) DAPI+ overview images of representative sections from the 2-, 4-, 8-, and 40-hour EdU groups, corresponding to those shown in Figure 1 and panel (B), to visualize the relative abundance of EdU-labeled nuclei (magenta) across time points.

**<u>SUPPLEMENTAL FIGURE S2 Validation of electroporation efficiency in the developing little skate retina.</u>**

(A–B) Whole-mount views of retinas electroporated with control constructs showing robust reporter expression driven by the ubiquitous CAG promoter. (A) CAG::mCherry and (B) CAG::GFP demonstrate efficient transfection across the retinal surface. Scale bar in A = 200 µm and applies to B. (C–E) Cryosections of electroporated retinas. (C) CAG::GFP and (D) CAG::mCherry label retinal cells spanning the neuroepithelium. (E) Merge with DAPI (blue) highlights nuclear organization and confirms widespread distribution of electroporated cells. Dashed line indicates the basal boundary of the retina. Scale bar in C = 50 µm and applies to D and E.

**<u>SUPPLEMENTAL FIGURE S3 Comparative expression and genomic context of cone phototransduction genes absent in the skate genome.</u>**

(A) Table summarizing TBLASTN hits recovered from the *Leucoraja erinacea* genome using human and zebrafish NRL protein sequences as queries. Hits are ranked by bit score and listed with corresponding locus, chromosome, gene ID, bit score, E value, and percent identity. Top-scoring hits correspond to other Maf family members rather than a distinct nrl ortholog. (B) Top-aligning genomic region in the *Leucoraja erinacea* genome for each pseudogene recovered by TBLASTN using elephant shark protein sequences as queries. Shown are aligning regions for pde6c, arr3, gnat2, and cngb3. Yellow highlights indicate nonsense mutations occurring within otherwise conserved regions, consistent with pseudogenization. (C) NCBI Genome Data Viewer (GDV) browser tracks for representative cone-associated loci shown in the order ghost shark, little skate, catshark, and devil ray. In the ghost shark track, the target gene is labeled at its annotated position. In the other species, the target gene is labeled where present; where no annotated target gene was present, two reference genes marking the corresponding conserved syntenic region are shown where available, and the genomic interval searched by targeted TBLASTN is indicated with a red line. In the little skate, this targeted interval corresponds to the region examined for recovery of the expected ortholog. In the devil ray gngt2 region, only a single reference gene (tut1) was available without identified syntenic partner genes, so only the position of tut1 and the searched genomic interval are shown. (D) TBLASTN results for vertebrate opsin queries (Ghost shark (Gshark) LWS1, LWS2, and RH2) against the little skate genome. Top hits correspond primarily to rhodopsin and other GPCR-related genes (e.g., pinopsin and Valopa), with no clear matches to cone opsin genes, consistent with the absence of canonical cone opsins in the little skate genome. (E) RNA-seq expression summary for selected transcription factors and phototransduction genes across developmental stages (embryo, hatchling, and adult). Genes are categorized by pathway and photoreceptor association (cone, rod, or shared), showing robust expression of rod phototransduction components and transcription factors across stages.

**<u>SUPPLEMENTAL FIGURE S4 Analysis of *Onecut1* splice isoform usage and spacer region conservation across development and species.</u>**

(A) Table of the number and frequency of Onecut1 spacer and canonical isoform reads at three developmental stages in little skate. (B) Alignment of bulk RNA-seq reads containing the truncation query sequence (5′-ACAAGGAGAT-3′) to the *onecut1* cDNA library-derived sequence and the reference genomic sequence. All query-matching reads aligned to the canonical reference continuation (green), with occasional single nucleotide variants (grey). No reads matched the cDNA library-derived continuation (red), suggesting it is a technical artifact. (C) IGV screenshot comparing RNA-seq coverage and splice junctions across the Onecut1 locus in little skate and mouse embryonic retina. The little skate embryo (S29) shows abundant inclusion of a spacer exon (arrowhead), supported by junction-spanning reads. In contrast, wild-type C57BL/6N mouse retina at E14.5 (Brooks et al., 2019; panel D) and C57BL/6 mouse retina at E14.5 (Aldiri et al., 2017; panel E) display canonical splicing of *Onecut1* exons 1 and 2 with minimal inclusion of the analogous spacer region. Both mouse datasets show low-level read coverage in a region encoding codons analogous to the rat liver onecut1 spacer exon, but expression in this region is less than that of the embryonic skate spacer exon, and its functional significance is unclear. (F) Alignment of spacer region protein sequences from *onecut1* transcripts across four myliobatiform species (*Mobula hypostoma*, *Hypanus sabinus*, *Hemitrygon akajei*, *Narcine bancroftii*) compared to *Leucoraja erinacea* (little skate). Sequences were obtained from NCBI standard reference genomes: *Mobula hypostoma*: sMobHyp1.1, *Hypanus sabinus*: sHypSab1.hap1, *Hemitrygon akajei*: sHemAka1.3, *Narcine bancroftii*: sNarBan1.hap1. Red indicates amino acid substitutions relative to little skate.

