## Supplemental material for "Developmental analysis of the cone photoreceptor-less little skate retina reveals distinct Onecut1 isoforms"

Supplemental Figure 1

**A**

| Time Elapsed After Edu Pulse | # of Samples | # of Cells | # Edu+ Cells |
|---|---|---|---|
| 4 Hours | 4 | 87 | 1 |
| 8 Hours | 4 | 56 | 43 |
| | Sample # | # of Cells | # Edu+ Cells |
| 4 Hours | 1 | 33 | 0 |
| | 2 | 16 | 0 |
| | 3 | 16 | 0 |
| | 4 | 22 | 1 |
| 8 Hours | 1 | 11 | 7 |
| | 2 | 14 | 12 |
| | 3 | 16 | 12 |
| | 4 | 15 | 12 |

**B**

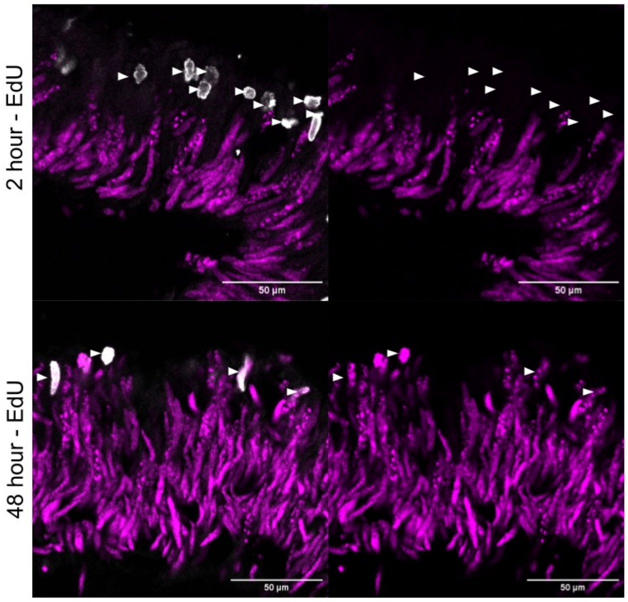


**C**

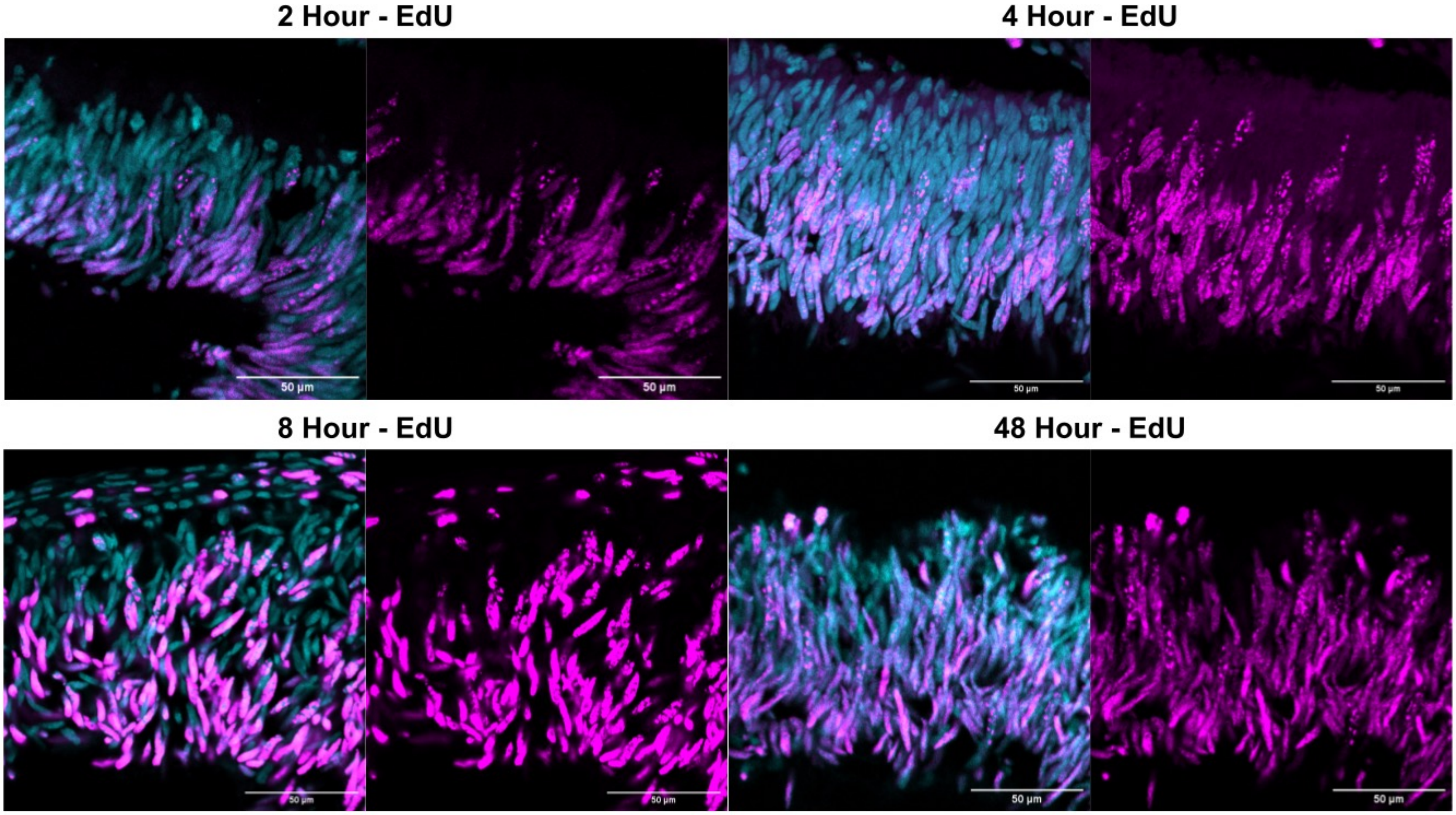

Supplemental Figure 2

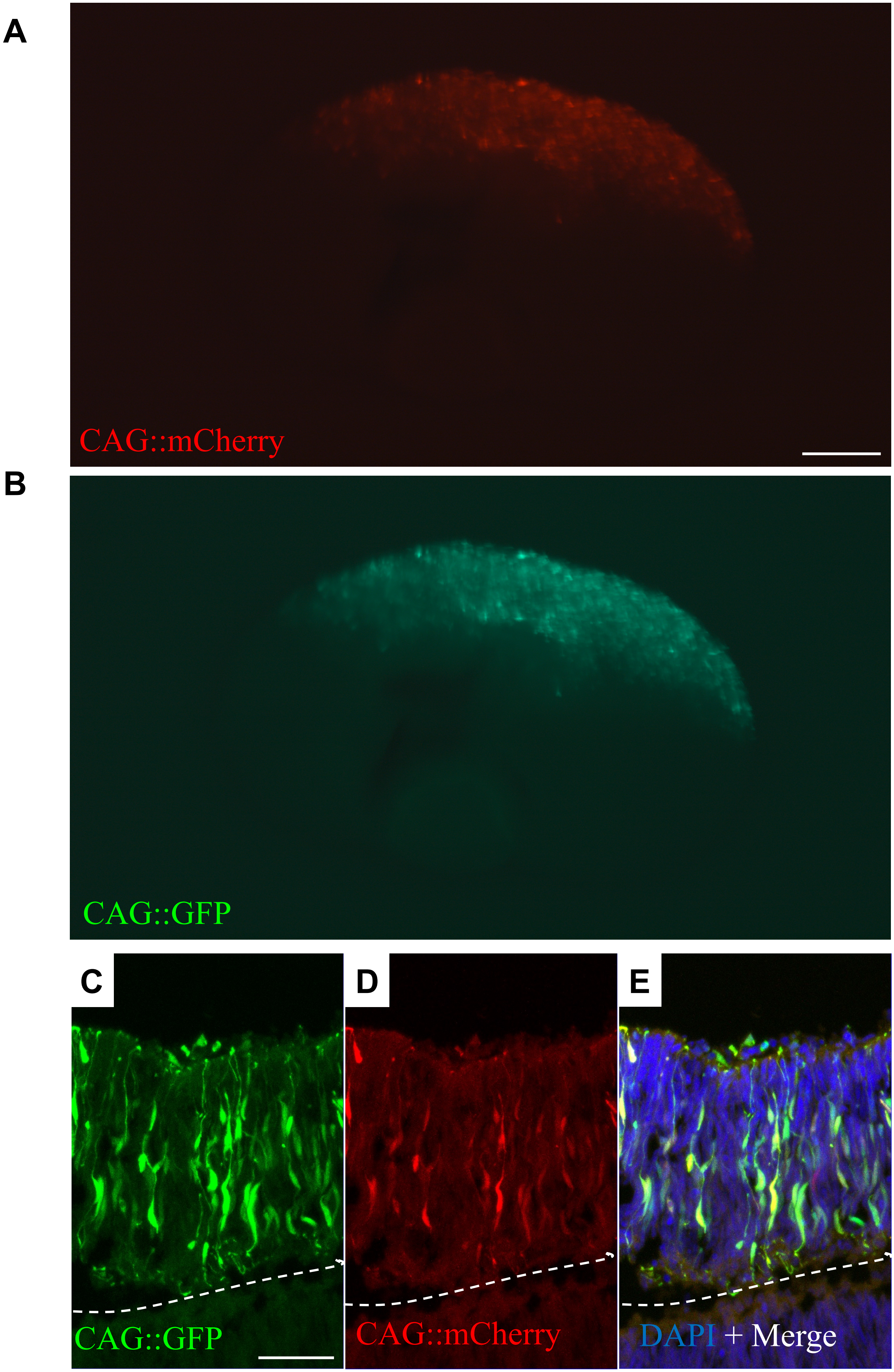

### Supplemental Figure 3

**A**

Nrl Homology Search

| Rank | Query | Locus | Chromosome | Gene ID | Score (bits) | E-value | Identity % |
|---|---|---|---|---|---|---|---|
| 1 | Human NRL | NC_073397.1 | chr21 | mafb | 130 | 1E-31 | 39% (113/292) |
| 2 | Human NRL | NC_073380.1 | chr4 | mafaa | 129 | 4E-31 | 44% (96/216) |
| 3 | Human NRL | NC_073393.1 | chr17 | c-maf | 90.9 | 5E-18 | 61% (62/102) |
| 4 | Human NRL | NC_073399.1 | chr23 | mafg | 75.5 | 6E-13 | 68% (34/50) |
| 5 | Human NRL | NC_073396.1 | chr20 | mafk | 72.8 | 5E-12 | 66% (33/50) |
| 1 | Zebrafish NRL | NC_073393.1 | chr17 | c-maf | 136 | 1E-32 | 73% (73/100) |
| 2 | Zebrafish NRL | NC_073380.1 | chr4 | mafaa | 133 | 1E-31 | 72% (73/101) |
| 3 | Zebrafish NRL | NC_073397.1 | chr21 | mafb | 130 | 1E-30 | 70% (71/102) |
| 4 | Zebrafish NRL | NC_073399.1 | chr23 | mafg | 91.7 | 8E-18 | 53% (49/93) |
| 5 | Zebrafish NRL | NC_073413.1 | chr37 | mafk-like | 87.8 | 1E-16 | 53% (48/91) |

**B**

**Pde6c**

```
Score  =  90.1 bits  (222),   Expect  = 3e-28,  Method:  Compositional  matrix
adjust.
 Identities = 43/48 (90%), Positives = 44/48 (92%), Gaps = 0/48 (0%)
 Frame = -1

Query  249    QVLLWSANKVFEELTDIERQFHKALYTVRIYLNVERYSVALLDMTKQK  296
              Q+LLWS NKVFEEL DI  QFHKALYTVRIYLNVERYSVALLDMTKQK
Sbjct  18153  QILLWSDNKVFEELADI*HQFHKALYTVRIYLNVERYSVALLDMTKQK  18010
```

**Arr3**

```
Score  =  55.8 bits  (133),   Expect  = 3e-09,  Method:  Compositional  matrix
adjust.
 Identities = 27/39 (69%), Positives = 33/39 (85%), Gaps = 0/39 (0%)
 Frame = +2

Query  177    RKVQFAPEESKVKPKVETTRQFLMSDKPLNLQVSLEKEL  215
              RKVQ A EES ++P+VETTRQFLMSDKPL L+ +L KE+
Sbjct  65591  RKVQ*ALEESLLQPRVETTRQFLMSDKPLQLEATLGKEV  65707
```

**Gnat2**

```
Score  =  79.3 bits  (194),   Expect  = 8e-18,  Method:  Compositional  matrix
adjust.
Identities = 35/43 (81%), Positives = 39/43 (91%), Gaps = 0/43 (0%)
Frame = +2

Query  227    DVGGQRSERKKWIHCFEGVTCIIFCGALSAYDMVLVEDDDVNR  269
              DVGGQ S RK+W HCFEGVTC+IFCGALSAY+MVLVEDDD+ R
Sbjct  13517  DVGGQVS*RKRWFHCFEGVTCMIFCGALSAYNMVLVEDDDMVR  13645
```

**Cngb3**

```
Score = 64.7 bits (156),  Expect(2) = 3e-18, Method: Compositional matrix
adjust.
Identities = 28/53 (53%), Positives = 41/53 (77%), Gaps = 0/53 (0%)
Frame = +1

Query  474     AATSAQNYFRSSMDNTVHYMNINSIPKIVHNRVRTWYEYTWKSQGILDESELL  526
               AAT+ Q+Y+++S+DNTV +MNI ++ + V NR+R W EYTW+SQG L +  LL
Sbjct  414226  AATAGQSYYQTSLDNTVAFMNIYAVSRNVQNRIRKWDEYTWESQGQLGK*ALL  414384
```

C

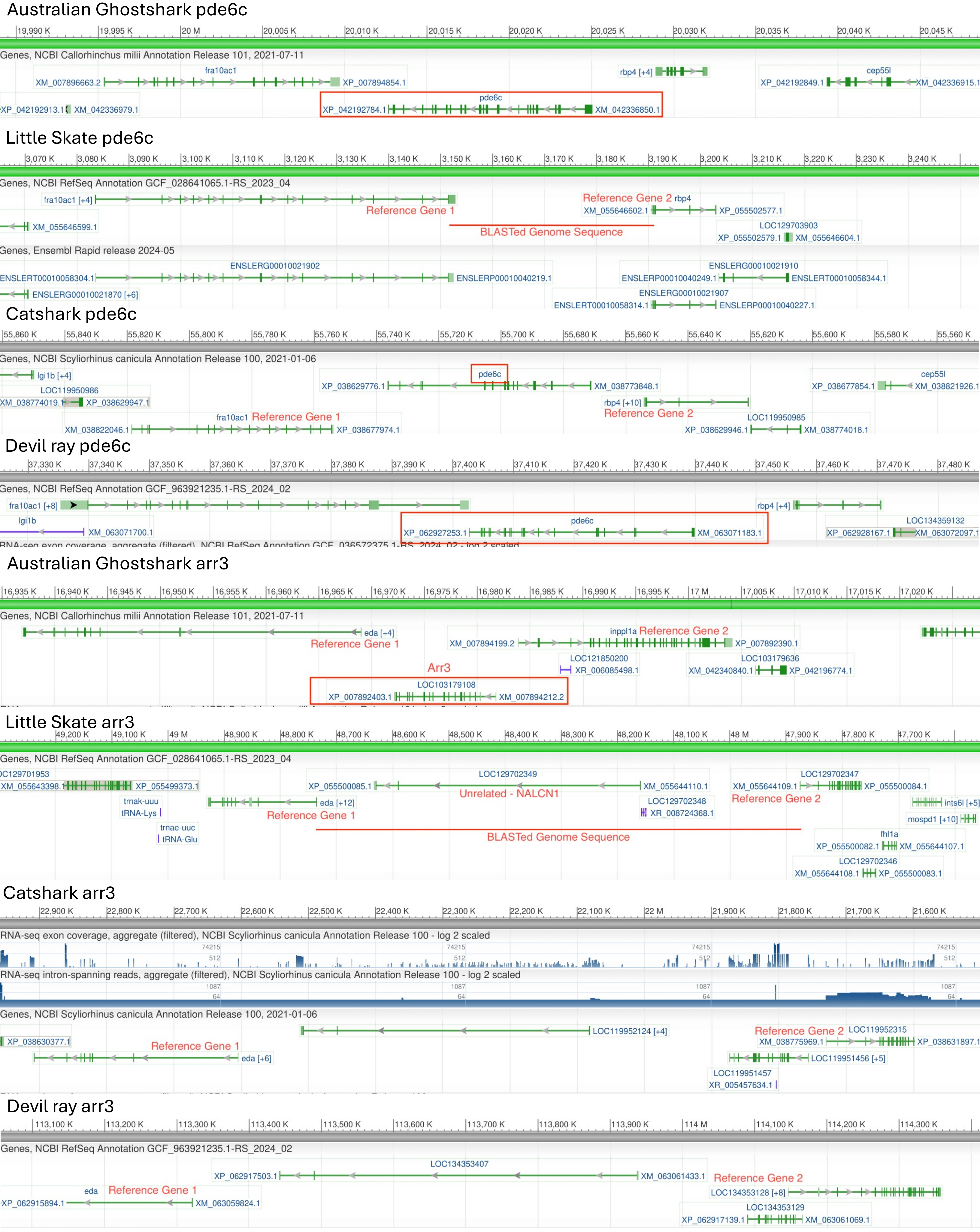

Australian Ghostshark pde6c
Genes, NCBI Callorhinchus milii Annotation Release 101, 2021-07-11
fra10ac1
XM_007896663.2
XP_007894854.1
rbp4 [+4]
cep55l
XP_042192849.1
XM_042336915.1
pde6c
XP_042192913.1
XM_042336979.1
XP_042192784.1
XM_042336850.1
Little Skate pde6c
Genes, NCBI RefSeq Annotation GCF_028641065.1-RS_2023_04
fra10ac1 [+4]
Reference Gene 1
Reference Gene 2
rbp4
XM_055646602.1
XP_055502577.1
XM_055646599.1
BLASTed Genome Sequence
LOC129703903
XP_055502579.1
XM_055646604.1
Genes, Ensembl Rapid release 2024-05
ENSLERG00010021902
ENSLERT00010058304.1
ENSLERP00010040219.1
ENSLERG00010021910
ENSLERP00010040249.1
ENSLERT00010058344.1
ENSLERG00010021870 [+6]
ENSLERG00010021907
ENSLERT00010058314.1
ENSLERP00010040227.1
Catshark pde6c
Genes, NCBI Scyliorhinus canicula Annotation Release 100, 2021-01-06
lgi1b [+4]
pde6c
XP_038629776.1
XM_038773848.1
cep55l
XP_038677854.1
XM_038821926.1
LOC119950986
XM_038774019.1
XP_038629947.1
rbp4 [+10]
fra10ac1
Reference Gene 1
Reference Gene 2
LOC119950985
XM_038822046.1
XP_038677974.1
XP_038629946.1
XM_038774018.1
Devil ray pde6c
Genes, NCBI RefSeq Annotation GCF_963921235.1-RS_2024_02
fra10ac1 [+8]
rbp4 [+4]
lgi1b
pde6c
LOC134359132
XM_063071700.1
XP_062927253.1
XM_063071183.1
XP_062928167.1
XM_063072097.1
Australian Ghostshark arr3
Genes, NCBI Callorhinchus milii Annotation Release 101, 2021-07-11
eda [+4]
Reference Gene 1
inppl1a
Reference Gene 2
XM_007894199.2
XP_007892390.1
Arr3
LOC121850200
XR_006085498.1
LOC103179636
XM_042340840.1
XP_042196774.1
LOC103179108
XP_007892403.1
XM_007894212.2
Little Skate arr3
Genes, NCBI RefSeq Annotation GCF_028641065.1-RS_2023_04
LOC129701953
XM_055643398.1
XP_055499373.1
LOC129702349
LOC129702347
XP_055500085.1
XM_055644110.1
XM_055644109.1
XP_055500084.1
trnak-uuu
tRNA-Lys
eda [+12]
Unrelated - NALCN1
LOC129702348
XR_008724368.1
Reference Gene 2
ints6l [+5]
Reference Gene 1
mospd1 [+10]
trnae-uuc
tRNA-Glu
BLASTed Genome Sequence
fhl1a
XP_055500082.1
XM_055644107.1
LOC129702346
XM_055644108.1
XP_055500083.1
Catshark arr3
RNA-seq exon coverage, aggregate (filtered), NCBI Scyliorhinus canicula Annotation Release 100 - log 2 scaled
RNA-seq intron-spanning reads, aggregate (filtered), NCBI Scyliorhinus canicula Annotation Release 100 - log 2 scaled
Genes, NCBI Scyliorhinus canicula Annotation Release 100, 2021-01-06
LOC119952124 [+4]
Reference Gene 2
LOC119952315
XP_038630377.1
Reference Gene 1
XM_038775969.1
XP_038631897.1
eda [+6]
LOC119951456 [+5]
LOC119951457
XR_005457634.1
Devil ray arr3
Genes, NCBI RefSeq Annotation GCF_963921235.1-RS_2024_02
LOC134353407
XP_062917503.1
XM_063061433.1
Reference Gene 2
eda
Reference Gene 1
LOC134353128 [+8]
XP_062915894.1
XM_063059824.1
LOC134353129
XP_062917139.1
XM_063061069.1

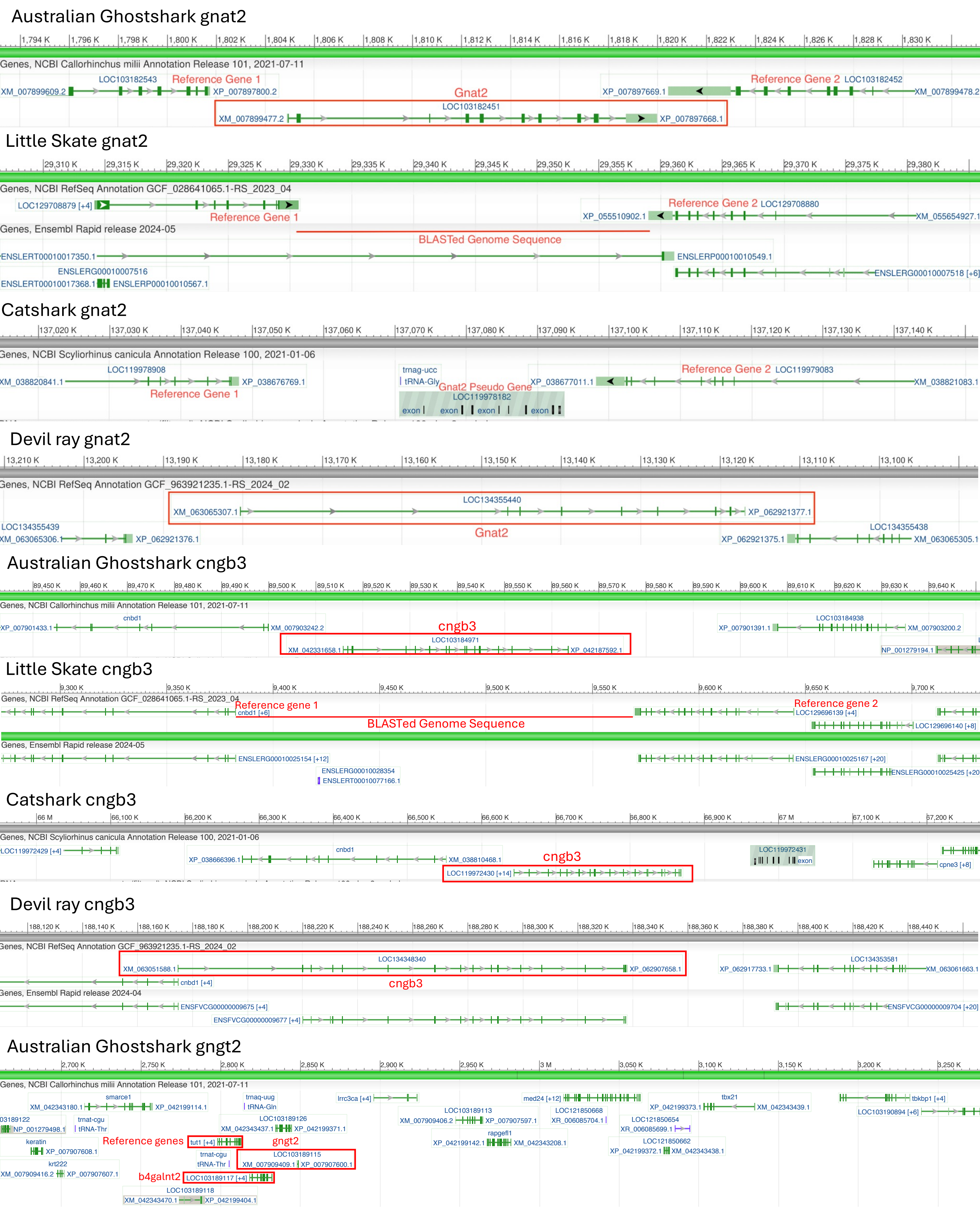

Australian Ghostshark gnat2
Genes, NCBI Callorhinchus milii Annotation Release 101, 2021-07-11
LOC103182543
Reference Gene 1
XM_007899609.2
XP_007897800.2
Gnat2
LOC103182451
XM_007899477.2
XP_007897668.1
Reference Gene 2
LOC103182452
XP_007897669.1
XM_007899478.2
Little Skate gnat2
Genes, NCBI RefSeq Annotation GCF_028641065.1-RS_2023_04
LOC129708879 [+4]
Reference Gene 1
Reference Gene 2
LOC129708880
XP_055510902.1
XM_055654927.1
Genes, Ensembl Rapid release 2024-05
BLASTed Genome Sequence
ENSLERT00010017350.1
ENSLERP00010010549.1
ENSLERG00010007516
ENSLERT00010017368.1
ENSLERP00010010567.1
ENSLERG00010007518 [+6]
Catshark gnat2
Genes, NCBI Scyliorhinus canicula Annotation Release 100, 2021-01-06
LOC119978908
XM_038820841.1
XP_038676769.1
Reference Gene 1
trnag-ucc
tRNA-Gly
Gnat2 Pseudo Gene
LOC119978182
exon
XP_038677011.1
Reference Gene 2
LOC119979083
XM_038821083.1
Devil ray gnat2
Genes, NCBI RefSeq Annotation GCF_963921235.1-RS_2024_02
LOC134355440
XM_063065307.1
XP_062921377.1
Gnat2
LOC134355439
XM_063065306.1
XP_062921376.1
LOC134355438
XP_062921375.1
XM_063065305.1
Australian Ghostshark cngb3
Genes, NCBI Callorhinchus milii Annotation Release 101, 2021-07-11
cnbd1
XP_007901433.1
XM_007903242.2
cngb3
LOC103184971
XM_042331658.1
XP_042187592.1
LOC103184938
XP_007901391.1
XM_007903200.2
NP_001279194.1
Little Skate cngb3
Genes, NCBI RefSeq Annotation GCF_028641065.1-RS_2023_04
Reference gene 1
cnbd1 [+6]
BLASTed Genome Sequence
Reference gene 2
LOC129696139 [+4]
LOC129696140 [+8]
Genes, Ensembl Rapid release 2024-05
ENSLERG00010025154 [+12]
ENSLERG00010028354
ENSLERT00010077166.1
ENSLERG00010025167 [+20]
ENSLERG00010025425 [+20]
Catshark cngb3
Genes, NCBI Scyliorhinus canicula Annotation Release 100, 2021-01-06
LOC119972429 [+4]
cnbd1
XP_038666396.1
XM_038810468.1
cngb3
LOC119972430 [+14]
LOC119972431
exon
cpne3 [+8]
Devil ray cngb3
Genes, NCBI RefSeq Annotation GCF_963921235.1-RS_2024_02
LOC134348340
XM_063051588.1
XP_062907658.1
cngb3
cnbd1 [+4]
LOC134353581
XP_062917733.1
XM_063061663.1
Genes, Ensembl Rapid release 2024-04
ENSFVCG00000009675 [+4]
ENSFVCG00000009677 [+4]
ENSFVCG00000009704 [+20]
Australian Ghostshark gngt2
Genes, NCBI Callorhinchus milii Annotation Release 101, 2021-07-11
smarce1
XM_042343180.1
XP_042199114.1
trnaq-uug
tRNA-Gln
lrrc3ca [+4]
med24 [+12]
tbx21
tbkbp1 [+4]
LOC103189126
XM_042343437.1
XP_042199371.1
LOC103189113
XM_007909406.2
XP_007907597.1
LOC121850668
XR_006085704.1
XP_042199373.1
XM_042343439.1
LOC103190894 [+6]
trnat-cgu
tRNA-Thr
NP_001279498.1
LOC121850654
XR_006085699.1
rapgef1
XP_042199142.1
XM_042343208.1
keratin
XP_007907608.1
Reference genes
tut1 [+4]
gngt2
LOC103189115
XM_007909409.1
XP_007907600.1
LOC121850662
XP_042199372.1
XM_042343438.1
krt222
XM_007909416.2
XP_007907607.1
b4galnt2
LOC103189117 [+4]
LOC103189118
XM_042343470.1
XP_042199404.1

#### Little Skate gngt2

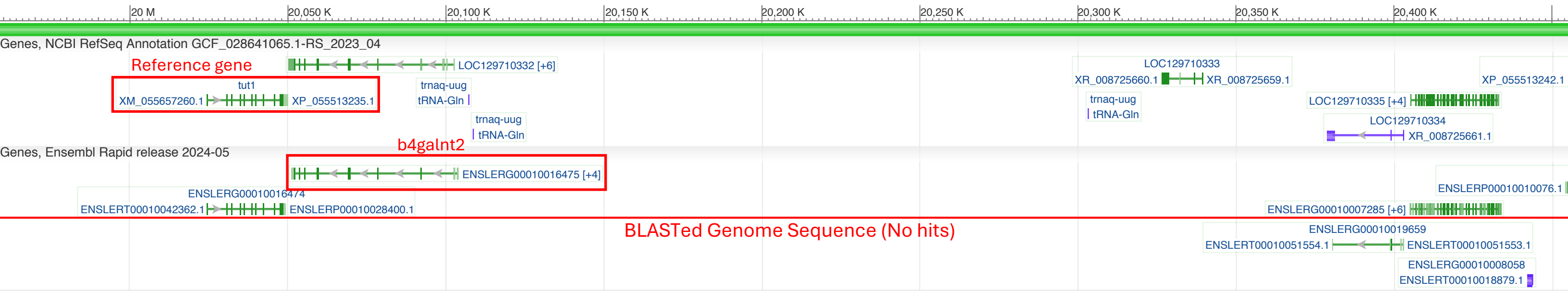


#### Catshark gngt2

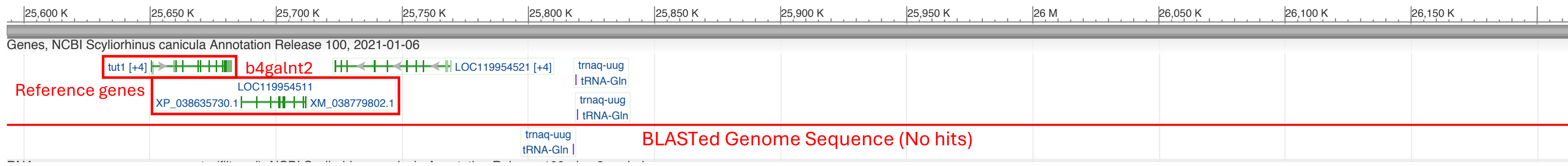


#### Devil ray gngt2

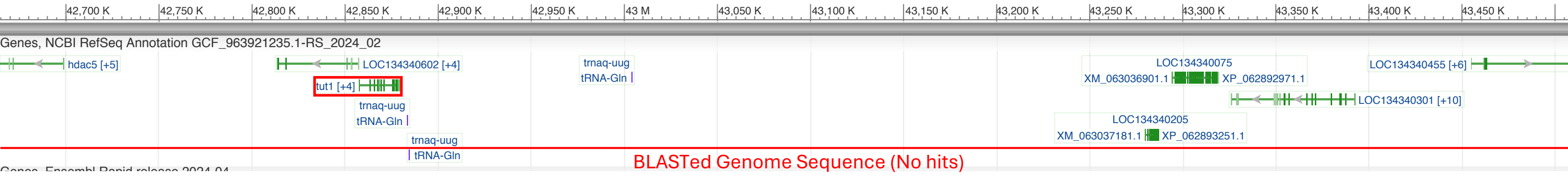

**D**

TBLASTN hits for opsin genes

| Query | Rank | Accession | Chromosome | Score_bits | E_value | Gene_at_locus |
|---|---|---|---|---|---|---|
| **Gshark LWS1** | 1 | NC_073404.1 | chr28 | 115 | 4E-26 | pinopsin |
| **Gshark LWS1** | 2 | NC_073392.1 | chr16 | 103 | 4E-22 | rhodopsin |
| **Gshark LWS1** | 3 | NC_073385.1 | chr9 | 90.5 | 1E-17 | G-protein coupled receptor 135 |
| **Gshark LWS1** | 4 | NC_073391.1 | chr15 | 84.7 | 9E-16 | Valopa |
| **Gshark LWS1** | 5 | NC_073414.1 | chr38 | 73.2 | 6E-12 | adenosine receptor A1-like |
| **Gshark LWS2** | 1 | NC_073404.1 | chr28 | 114 | 8E-26 | pinopsin |
| **Gshark LWS2** | 2 | NC_073392.1 | chr16 | 104 | 3E-22 | rhodopsin |
| **Gshark LWS2** | 3 | NC_073391.1 | chr15 | 92.4 | 3E-18 | Valopa |
| **Gshark LWS2** | 4 | NC_073385.1 | chr9 | 88.6 | 5E-17 | G-protein coupled receptor 135 |
| **Gshark LWS2** | 5 | NC_073387.1 | chr11 | 71.2 | 2E-11 | rhodopsin-like |
| **Gshark RH2** | 1 | NC_073392.1 | chr16 | 165 | 3E-43 | rhodopsin |
| **Gshark RH2** | 2 | NC_073404.1 | chr28 | 81.6 | 7E-15 | pinopsin |
| **Gshark RH2** | 3 | NC_073391.1 | chr15 | 75.5 | 8E-13 | Valopa |
| **Gshark RH2** | 4 | NC_073387.1 | chr11 | 70.9 | 3E-11 | neuropeptide Y receptor type 6-like |
| **Gshark RH2** | 5 | NC_073393.1 | chr17 | 58.2 | 3E-07 | thyrotropin-releasing hormone receptor-like |
| **Gshark RH2** | 6 | NC_073397.1 | chr21 | 56.6 | 9E-07 | cadherin-4-like |
| **Gshark RH2** | 7 | NC_073377.1 | chr1 | 56.2 | 1E-06 | retinal pigment epithelium-derived rhodopsin homolog |
| **Gshark RH2** | 8 | NC_073396.1 | chr20 | 56.2 | 1E-06 | urotensin-2 receptor 5 |
| **Gshark RH2** | 9 | NC_073390.1 | chr14 | 52.8 | 1E-05 | angiotensin II receptor, type 1a |
| **Gshark RH2** | 10 | NC_073411.1 | chr35 | 52.8 | 2E-05 | adrenoceptor alpha 1Aa |
| **Gshark RH2** | 11 | NC_073382.1 | chr6 | 51.2 | 5E-05 | melatonin receptor type 1B-like |
| **Gshark RH2** | 12 | NC_073384.1 | chr8 | 50.8 | 7E-05 | opsin 3 |
| **Gshark RH2** | 13 | NC_073381.1 | chr5 | 49.7 | 2E-04 | opsin 6, group member a |
| **Gshark RH2** | 14 | NC_073400.1 | chr24 | 49.7 | 2E-04 | G-protein coupled receptor moody-like |

**E**

gene_expression_table

| Gene | Pathway | Group | Embryo_Avg | Hatchling_Avg | Adult_Avg | Embryo_SE | Hatchling_SE | Adult_SE | Embryo_pSE | Hatchling_pSE | Adult_pSE |
|---|---|---|---|---|---|---|---|---|---|---|---|
| **otx2b** | Shared | Transcription Factors | 14.07229983 | 30.65873673 | 14.47604855 | 0.401362029 | 0.443247003 | 0.056746807 | 2.85214239213662 | 1.4457445096434 | 0.392004812666921 |
| **otx5 (crx)** | Shared | Transcription Factors | 58.86898384 | 1310.843866 | 758.8196186 | 2.119836098 | 11.96447406 | 91.72577 | 3.60093883013422 | 0.912730674516503 | 12.0879544692362 |
| **rxrga** | Cone | Transcription Factors | 7.207887584 | 17.41604708 | 10.78790285 | 0.522505019 | 1.611727353 | 0.93230113 | 7.24907280962389 | 9.25426616956527 | 8.64209794028688 |
| **mafb** | Rod | Transcription Factors | 1.87241533 | 34.34727073 | 29.31883008 | 0.354528524 | 1.159286504 | 2.326405425 | 18.9342886869015 | 3.37519249524371 | 7.93485080629793 |
| **mafaa** | Rod | Transcription Factors | 0.09498602 | 3.888494566 | 2.132791064 | 0.047550184 | 0.555288455 | 0.55982215 | 50.060192015625 | 14.28029396917 | 26.2483353127927 |
| **neurod1** | Shared | Transcription Factors | 157.0264333 | 416.7631267 | 301.0623991 | 3.653252745 | 17.22425214 | 6.774600329 | 2.32652087182063 | 4.13286373878239 | 2.25023129731646 |
| **prdm1a** | Cone | Transcription Factors | 0.194364992 | 0.232085411 | 0.183415542 | 0.013198549 | 0.075337544 | 0.05355762 | 6.79059992449669 | 32.4611287178236 | 29.2001536053035 |
| **nr2e3** | Rod | Transcription Factors | 4.557174404 | 230.4304814 | 89.60939484 | 0.759805694 | 30.4818186 | 23.53089983 | 16.6727368022846 | 13.2282059277944 | 26.2594116074716 |
| **rorb** | Cone | Transcription Factors | 5.910970358 | 29.88027479 | 13.33382639 | 0.342209403 | 5.538505926 | 1.069046806 | 5.78939467251512 | 18.5356592766462 | 8.01755456184547 |
| **thrb** | Cone | Transcription Factors | 0.214125347 | 1.341506528 | 0.839320464 | 0.024968371 | 0.191496543 | 0.12769306 | 11.6606330590091 | 14.2747380652329 | 15.2138623418575 |
| **onecut1** | Cone | Transcription Factors | 46.09665192 | 25.28489608 | 6.67183396 | 0.911631676 | 1.870493337 | 2.441198609 | 1.97765268849053 | 7.397670653191 | 36.5896187410515 |
| **hprt1** | Control | Transcription Factors | 35.8810947 | 39.289309 | 40.84258471 | 2.71107851 | 3.941891616 | 0.079328427 | 7.55572964723398 | 10.0329878950022 | 0.194229693255865 |
| **c-maf** | Rod | Transcription Factors | 1.272528761 | 3.755419671 | 1.979997174 | 0.370809954 | 0.840508241 | 0.128659631 | 29.1396128216862 | 22.3812067527512 | 6.49797043599215 |
| **olig2** | Shared | Transcription Factors | 114.3427784 | 4.372624635 | 3.306371727 | 3.62084196 | 0.791808903 | 0.20117602 | 3.16665556904117 | 18.1083209535547 | 6.0844949270884 |
| **rho** | Rod | Phototransduction | 5.38023505 | 76274.47255 | 98896.23224 | 0.727846322 | 10949.56454 | 9718.920852 | 13.5281510052242 | 14.3554772310254 | 9.82739244141704 |
| **gnat1** | Rod | Phototransduction | 1.203146007 | 5748.887097 | 8832.016929 | 0.739401092 | 917.5486687 | 1490.948286 | 61.4556411024186 | 15.9604572714398 | 16.8811755908717 |
| **gnb1** | Rod | Phototransduction | 267.0197585 | 3802.7411 | 4077.153284 | 9.339088073 | 516.1257172 | 437.1945492 | 3.49752697158551 | 13.5724653250783 | 10.7230344003912 |
| **gngt1** | Rod | Phototransduction | 0.093008304 | 48.48413648 | 61.53265209 | 0.011966519 | 6.309933067 | 6.457206279 | 12.866075915114 | 13.0144280688651 | 10.4939508694594 |
| **pde6a** | Rod | Phototransduction | 0.062088339 | 555.934554 | 639.8218037 | 0.036536081 | 64.37409644 | 101.2127708 | 58.8453187642852 | 11.5794378990877 | 15.8188999209937 |
| **pde6b** | Rod | Phototransduction | 0.165807563 | 561.8113208 | 725.4867427 | 0.02511377 | 90.53087725 | 79.1948522 | 15.146335634883 | 16.1141069783156 | 10.9160991564457 |
| **pde6g** | Rod | Phototransduction | 0.558822484 | 2154.577718 | 2906.913066 | 0.16469097 | 295.7253884 | 529.0327182 | 29.4710708168285 | 13.7254454053534 | 18.1991241632817 |
| **cnga1** | Rod | Phototransduction | 0.036906035 | 365.2258157 | 439.476991 | 0.021967143 | 45.37845978 | 73.9412061 | 59.5218180441221 | 12.4247678639656 | 16.8248185034106 |
| **cngb1** | Rod | Phototransduction | 1.619234013 | 398.2276531 | 424.8174827 | 0.174008222 | 18.10641434 | 87.74230132 | 10.7463294744908 | 4.5467496290252 | 20.6541173311274 |
| **grk1** | Rod | Phototransduction | 0.399683091 | 1667.216382 | 2143.048094 | 0.060094253 | 181.9452606 | 313.4659855 | 15.0354754437185 | 10.9131161716236 | 14.6271092271623 |
| **sag** | Rod | Phototransduction | 0.494604286 | 3761.796904 | 4684.050902 | 0.074623648 | 595.6886241 | 787.0162554 | 15.0875457638068 | 15.8352149066472 | 16.8020431858236 |
| **cnga3a** | Cone | Phototransduction | 1.987574701 | 2.732732123 | 1.24936353 | 0.507781964 | 0.360347233 | 0.023992258 | 25.5478178377156 | 13.1863357541393 | 1.92035844042926 |
| **gnb3** | Cone | Phototransduction | 0.118131557 | 531.2408687 | 482.6542519 | 0.059246275 | 66.75436325 | 82.68506098 | 50.1527927884672 | 12.5657431841332 | 17.1313234379486 |
| **rcvrn** | Shared | Phototransduction | 1.1013846 | 1938.001625 | 2542.467907 | 0.238667311 | 505.6739342 | 74.48550194 | 21.6697519649358 | 26.0925443857664 | 2.92965357536763 |
| **rgs9** | Shared | Phototransduction | 1.50486346 | 86.78020503 | 71.64673229 | 0.196533733 | 8.800747615 | 19.68788797 | 13.0599046507515 | 10.1414229339025 | 27.4791150143604 |
| **gnb5** | Shared | Phototransduction | 13.71751272 | 397.5372667 | 445.0907106 | 0.987619936 | 8.12057773 | 3.504378484 | 7.19970125896118 | 2.0427211258481 | 0.787340288292235 |
| **rgs9bp** | Shared | Phototransduction | 3.572336203 | 138.7897703 | 202.9205383 | 0.861631185 | 18.46832693 | 56.88115017 | 24.1195435154287 | 13.3066917612731 | 28.0312434840412 |
| **guca1a** | Shared | Phototransduction | 0.337419754 | 526.6679817 | 1054.038641 | 0.149600167 | 88.57240574 | 155.0336636 | 44.3365171204529 | 16.8175034020679 | 14.7085370089387 |
| **guca1b** | Shared | Phototransduction | 0.202442384 | 487.4632891 | 717.2258301 | 0.119911646 | 68.53901569 | 138.6243293 | 59.2324806844796 | 14.0603440756622 | 19.327849539478 |
| **gucy2f** | Shared | Phototransduction | 0.14143603 | 250.3826757 | 279.8655465 | 0.05674203 | 24.36346468 | 42.46881457 | 40.118511527791 | 9.73049138159682 | 15.1747205403149 |
| **gucy2d** | Shared | Phototransduction | 0.119607897 | 60.89217195 | 82.96940785 | 0.027800966 | 10.10882013 | 15.37538167 | 23.2434201230041 | 16.6011817385995 | 18.5313865295954 |
| **slc24a1** | Shared | Phototransduction | 0.169037688 | 295.2568955 | 489.1203107 | 0.086570283 | 75.01359608 | 109.4429001 | 51.2135985911024 | 25.4062131056987 | 22.3754560393069 |

### Supplemental Figure 4

**A**

| Developmental Time Point | # of Samples | # of Total Reads | # of OC1 Spacer Isoform Reads | # of OC1 Cannonical Isoform Reads | Frequency of Spacer Isoform | Frequency of Cannonical Isoform |
|---|---|---|---|---|---|---|
| Embryo | 3 | 75 | 52 | 24 | 68.42% | 31.58% |
| Hatchling | 3 | 25 | 5 | 20 | 20.00% | 80.00% |
| Adult | 2 | 9 | 2 | 7 | 22.22% | 77.78% |

**B**

Reference Seq: ...**AGCCACTACCACCACAAGGAGAT**GAGCGGCATCGGCCAGAGCCTGTCGCCGCTGAACGGTTCCCCGCTGACCA...
cDNA Seq: ...**AGCCACTACCACCACAAGGAGAT**CAACACCAGGGAGATCGCGCAGAGGATAACCACCGAACTGAAGCGATACA...
Bulk RNA-seq Reads:
...**AGCCACTACCACCACAAGGAGAT**GAGCGGCATCGGCCAGAGCCTGTCGCCGCTGAACGGTTCCCCGCTGACCA
...**AGCCACTACCACCACAAGGAGAT**GAGCGGCATGGGCCAGAGCCTGTCGCCGCTGAACGGTTCCCCGCTGACCA
...**AGCCACTACCACCACAAGGAGAT**GAGCGGCATCGGCCAGAGCCTGTCGCCGCTGAACGGTTCCCCGCTGACCA
...**AGCCACTACCACCACAAGGAGAT**GAGCGGCATCGGCCAGAGCCTGTCGCCGCTGAACGGTTCCCCGCTGACCA
...**AGCCACTACCACCACAAGGAGAT**GAGCGGCATCGGCCAGAGCCTGTCGCCGCTGAACGGTTCCCCGCTGAC
...**AGCCACTACCACCACAAGGAGAT**GAGCGGCATCGGCCAGAGCCTGTCGCCGCTGAACGGTTCCCCGCTGAC
...**AGCCACTACCACCACAAGGAGAT**GAGCGGCATCGGCCAGAGCCTGTCGCCGCTGAACGGTTCCCCGCTGAC
...**AGCCACTACCACCACAAGGAGAT**GAGCGGCATCGGCCAGAGCCTGTCGCCGCTGAACGGTTCCCCGCTGA
...**AGCCACTACCACCACAAGGAGAT**GAGCGGCATCGGCCAGAGCCTGTCGCCGCTGAACGGTTCCCCGCTG
...**AGCCACTACCACCACAAGGAGAT**GAGCGGCATCGGCCAGAGCCTGTCGCCGCTGAACGGTTC
...**AGCCACTACCACCACAAGGAGAT**GAGCGGCATCGGCCAGAGCCTGTCGCCGCTGAACGGTT
...**AGCCACTACCACCACAAGGAGAT**GAGCGGTATCGGCCAGAGCCTGTCGCCGCTGAACGGTT

**C**

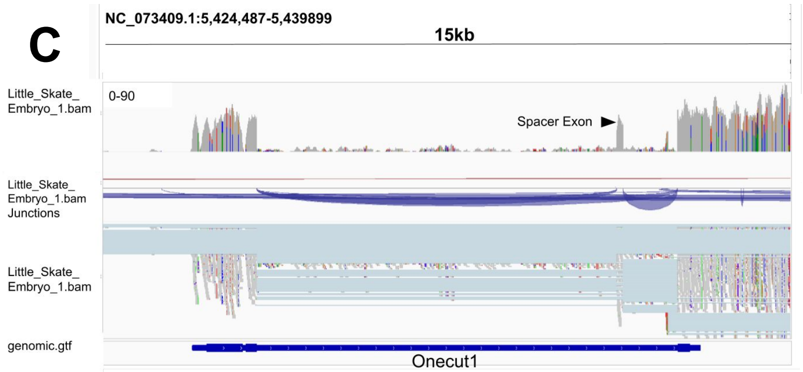


**D**

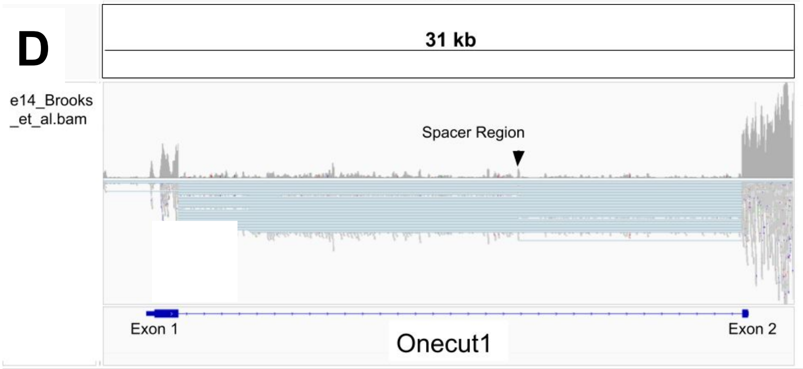


**E**

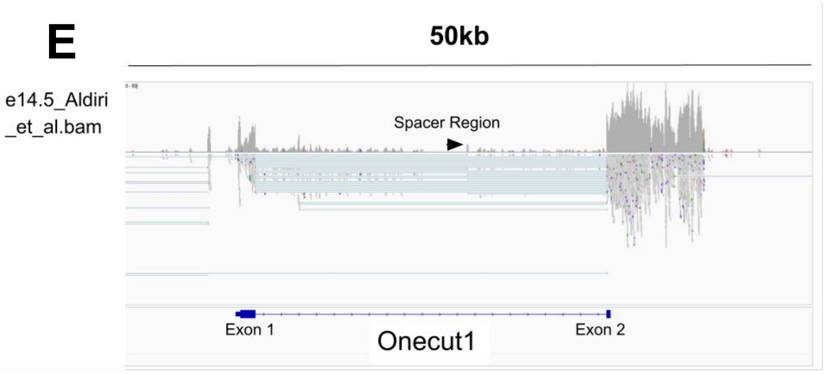


**F**

L. erinacea: GSQIGRQNEARCRAPTTWQLYSRELGEFLDRERTEEMDADLESDGSGQ
M. hypost.: GSQIGRQNDARYRAPTTWQLYSRELGEFLDRERTEEVDVDLEADGSRQ
H. sabinus: GSQIGRQNDARYRAPTTWQLYSRELGEFLDRERTEEVDVDLEADGSRQ
H. akajei: GSQIGRQNDARYRAPTTWQLYSRELGEFLDRERTEEVDVDLEADGSRQ
N. bancroftii: GSQFGRQNDARYRATTTWQLYSRELGEFLDREATEEVDADLESDGSRQ
